# Parkinson’s disease-related phosphorylation at Tyr39 rearranges α-synuclein amyloid fibril structure revealed by cryo-EM

**DOI:** 10.1101/2020.04.13.040261

**Authors:** Kun Zhao, Yeh-Jun Lim, Zhenying Liu, Houfang Long, Yunpeng Sun, Jin-Jian Hu, Chunyu Zhao, Youqi Tao, Xing Zhang, Dan Li, Yan-Mei Li, Cong Liu

## Abstract

Post-translational modifications (PTMs) of α-synuclein (α-syn), e.g. phosphorylation, play an important role in modulating α-syn pathology in Parkinson’s disease (PD) and α-synucleinopathies. Accumulation of phosphorylated α-syn fibrils in Lewy bodies and Lewy neurites is the histological hallmark of these diseases. However, it is unclear how phosphorylation relates to α-syn pathology. Here, by combining chemical synthesis and bacterial expression, we obtained homogeneous α-syn fibrils with site-specific phosphorylation at Y39, which exhibits enhanced neuronal pathology in rat primary cortical neurons. We determined the cryo-EM structure of pY39 α-syn fibril, which reveals a new fold of α-syn with pY39 in the center of the fibril core forming electrostatic interaction network with eight charged residues in the N-terminal region of α-syn. This structure composed of residues 1-100 represents the largest α-syn fibril core determined so far. This work provides structural understanding on the pathology of pY39 α-syn fibril, and highlights the importance of PTMs in defining the polymorphism and pathology of amyloid fibrils in neurodegenerative diseases.

## Introduction

Abnormal aggregation of α-synuclein (α-syn) into amyloid fibrils is closely associated with Parkinson’s disease (PD) and other synucleinopathies such as dementia with Lewy bodies and multiple system atrophy (MSA) (1, 2). Accumulation of α-syn fibrils in Lewy bodies (LBs) and Lewy neurites (LNs) is the histological hallmark of these diseases (3, 4). Moreover, α-syn fibril is the key pathological entity for the cell-to-cell transmission of α-syn pathology in brain or from other organs, such as gut, to the brain (5–8). A variety of post translational modifications (PTMs) of α-syn, e.g. phosphorylation, acetylation, ubiquitination and O-GlcNAcylation, have been identified in regulating α-syn biological function and pathological amyloid aggregation (9–12). Endogenous α-syn is generally acetylated at the N-terminus (12). α-Syn fibrils in LBs are commonly phosphorylated at S129 (pS129) and highly ubiquitinated in the N-terminus (3, 13). O-GlcNAcylation in the NAC region of α-syn (e.g. T72 and S87) can slow down or even eliminate α-syn fibrillation (9). Phosphorylation is a common PTM of α-syn. Besides pS129, phosphorylation has been reported on many other residues, including Y125, Y133 and Y135 in the C-terminus, and S87 in the NAC region (10, 14, 15). Recently, Y39 in the N-terminus has been identified to be phosphorylated by the c-Abl protein tyrosine kinase (16, 17). Phosphorylated Y39 (pY39) α-syn is elevated and accumulates in human brain tissues and Lewy bodies of sporadic PD patients (17). pY39 can decrease the lipid binding of α-syn, accelerate α-syn aggregation and neuropathology in transgenic mouse PD model (16–19). α-Syn phosphorylation by c-Abl protects α-syn against degradation via the autophagy and proteasome pathways in cortical neurons (16). However, the molecular mechanism of pY39 α-syn pathology is unknown.

In this work, we determined the near-atomic structure of full-length pY39 α-syn fibril by cryo-electron microscopy (cryo-EM). The structure comprises residues 1-100 covering the entire N-terminus, the NAC region and a short segment of the C-terminus. Strikingly, the structure shows that pY39 attracts the lysine residues in α-syn N-terminus to form a hydrophilic channel in the center of α-syn protofilament, which leads to complete rearrangement of α-syn fibril structure. In addition, we show that the pY39 α-syn fibril exhibits enhanced propagation in primary cortical neurons and is more resistant to protease digestion, compared to the non-phosphorylated WT fibril. This work provides structural indication for the pathology of pY39 α-syn fibril and suggests that PTM may alter α-syn fibril structure and affect PD pathology.

## Results

### Phosphorylation at Y39 alters α-syn fibril morphology

To investigate the influence of phosphorylation at Y39 on α-syn aggregation, we semi-synthesized full-length α-syn with site-specific phosphorylation at Y39 (Fig. 1*a* and *SI Appendix*, Fig. S1). The synthetic α-syn was characterized for the correct sequence and phosphorylation at Y39 by analytical reversed-phase high-performance liquid chromatography (RP-HPLC) and electrospray ionization mass spectrometry (ESI-MS) (*SI Appendix*, Fig. S2). pY39 α-syn amyloid fibril was grown in the buffer containing 50 mM Tris, 150 mM KCl, pH7.5 at 37 °C with agitation. The aggregation kinetics of pY39 α-syn was characterized and compared with that of non-phosphorylated WT α-syn by ThT kinetic assay. The result showed that the aggregation lag time of pY39 α-syn is ~50 hours similar to that of WT α-syn (Fig. 1*b*). Addition of preformed fibril seeds (PFFs) dramatically shortened the lag phase of both pY39 and WT α-syn aggregation (Fig. 1*b*). However, unlike the WT α-syn aggregation reaching plateau in ~100 hours, pY39 α-syn aggregation did not reach plateau during the monitored time (Fig. 1*b*). Consistently, negative-staining transmission electron microscopy (TEM) showed slower fibril formation of pY39 α-syn than that of WT α-syn (*SI Appendix*, Fig. S3*a*).

**Fig. 1.**
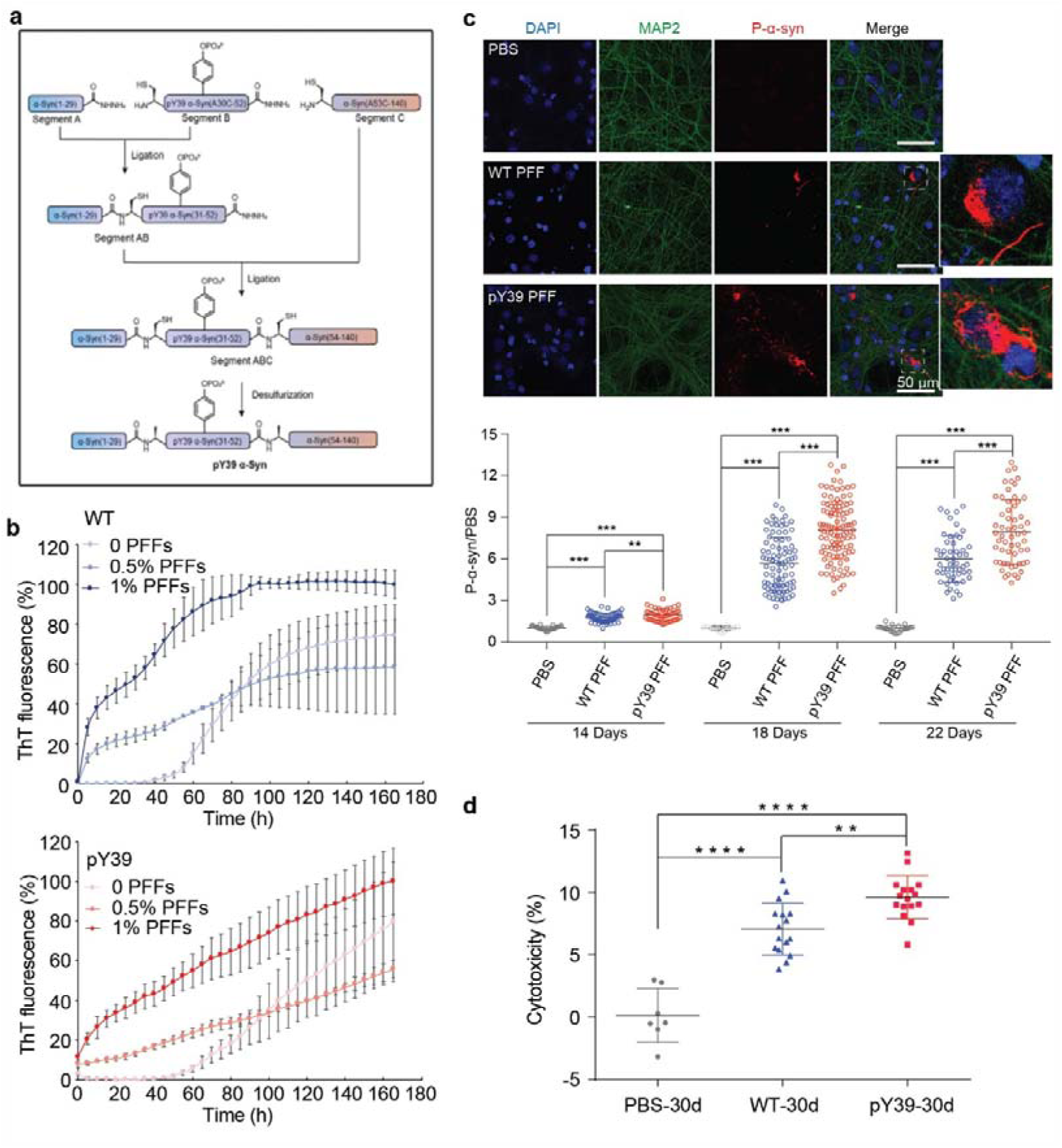
pY39 α-syn synthesis and pathology. (*a*) Workflow of the semi-synthesis of human pY39 α-syn. (*b*) ThT kinetic assays of the WT and pY39 aggregation with or without the addition of PFFs. Mol% of added PFFs are indicated. Data shown are mean ± s.d., n=3. (*c*) α-Syn PFFs induce endogenous α-syn aggregation in rat primary cortical neurons. Primary neurons were treated with 100 nM α-syn PFFs at DIV8 for 14 days. Fixed neurons were stained for DAPI (blue), MAP2 (green) and pS129 α-syn (red) and imaged by confocal microscopy. Measurement of the mean gray value of pS129 α-syn was shown in below. The images were analyzed by Image J. The mean gray values of pS129 α-syn were normalized to that of PBS. >10 images were randomly taken for each sample. >5 individual samples were analyzed for each time point. \*\**p*<0.01; \*\*\**p*<0.001 for one-way ANOVA followed by Tukey HSD post-hoc test. (*d*) Cytotoxicity of WT and pY39 α-syn PFFs to primary neurons at DIV8 for 30 days measured by LDH assay. **p<0.01; ****p<0.0001 for one-way ANOVA followed by Tukey HSD post-hoc test.

Furthermore, both negative-staining TEM and atomic force microscopy (AFM) revealed a mixture of three polymorphs of pY39 α-syn fibrils (*SI Appendix*, Fig. S3*b*). They include two types of left-handed twisted fibrils with a periodic spacing of ~122 nm and ~120 nm, respectively, and one type of straight fibril (*SI Appendix*, Fig. S3*c* and 3*d*). AFM measurement showed that ~80% of the total fibrils are twisted with a nearly equal proportion of the two types of twisted fibrils, while only ~20% are straight fibrils (*SI Appendix*, Fig. S3*e*). Notably, all the pY39 fibril polymorphs showed different morphologies from the non-phosphorylated WT α-syn fibril (referred to as WT fibril) formed under the same condition (*SI Appendix*, Fig. S4). The distinct morphologies of pY39 fibrils indicate that phosphorylation at Y39 may alter the structure of α-syn fibril.

### The pY39 α-syn fibril exhibits enhanced neuropathology in rat primary cortical neurons

It has been reported that phosphorylation at Y39 can exacerbate α-syn neuropathology (16,17). Thus, we sought to assess the biological relevance of the *in vitro* prepared pY39 α-syn fibril by testing its neuropathology. We firstly used a well-documented neuronal propagation assay to monitor the aggregation of endogenous α-syn induced by exogenous pY39 α-syn PFFs (6, 20, 21). We added PFFs into the culture medium of rat primary cortical neurons and detected the formation of pathological α-syn aggregation by the antibody for pS129 α-syn. The result showed that pY39 PFFs induced a significant elevation of pathological pS129 α-syn than the WT PFFs (Fig. 1*c* and *SI Appendix*, Fig. S5). To validate the aggregation property of the induced pS129 α-syn, we further assessed the immunoreactivity of PFF-treated neurons for Lewy-body markers: ubiquitin and P62, respectively. The result showed co-localization of pS129 α-syn, ubiquitin and P62, which confirms that the PFF-induced pS129 α-syn is pathological aggregation in Lewy bodies (*SI Appendix*, Fig. S6). Next, we used the lactate dehydrogenase (LDH) assay to evaluate the toxicity of pY39 fibril to neurons. The result showed that pY39 α-syn PFFs are more toxic to neurons than the WT PFFs (Fig. 1d). Together, these results indicate that the pY39 fibril prepared in this work exhibits consistent neuropathology to that of the pY39 α-syn aggregation *in vivo* (17).

### Cryo-EM structure determination of the pY39 α-syn amyloid fibril

To understand how phosphorylation at Y39 alters the structure and pathology of α-syn fibril, we set out to determine the atomic structure of pY39 α-syn fibrils by cryo-EM. We prepared cryo-EM grids with polydispersed pY39 fibrils and collected 3,177 micrographs. Consistent with the AFM data, 2D classification from helical image processing of 10,460 fibril segments revealed two major polymorphs of twisted fibrils (> 80% of the total): one consists of two protofilaments (termed twist-dimer); the other consists of three protofilaments (termed twist-trimer) (*SI Appendix*, Fig. S7). Besides, there was also a small species of straight fibril (< 20%), which consists of two protofilaments aligned in parallel (*SI Appendix*, Fig. S8).

We were able to reconstruct the 3D density maps for the two polymorphs of twisted fibrils (Fig. 2 and Table 1). The overall resolutions of the density maps are 3.22 Å for the twist-dimer fibril and 3.37 Å for the twist-trimer fibril (*SI Appendix*, Fig. S9). Both fibrils are reconstructed as left-handed helices according to AFM images (*SI Appendix*, Fig. S3*d*). The twist-dimer fibril spine features a width of ~18 nm and a half pitch (a pitch is termed as the length of a 360° helical turn of the whole fibril) of 124 nm. The two protofilaments intertwine along an approximate 2-fold screw axis with a helical twist of 179.65° (in the presence of an approximate 2-fold symmetry, the helical twist represents the relative turning angle of the two protofilaments) and rise of 2.41 Å (Fig. 2 and Table 1). The twist-trimer fibril spine features a width of ~18 nm and a half pitch of 125 nm. The helical twist of the twist-trimer fibril is −0.69° (in the absence of symmetry, the helical twist represents the turning angle of a protofilament) and rise of 4.80 Å (Fig. 2 and Table 1).

**Table 1.**
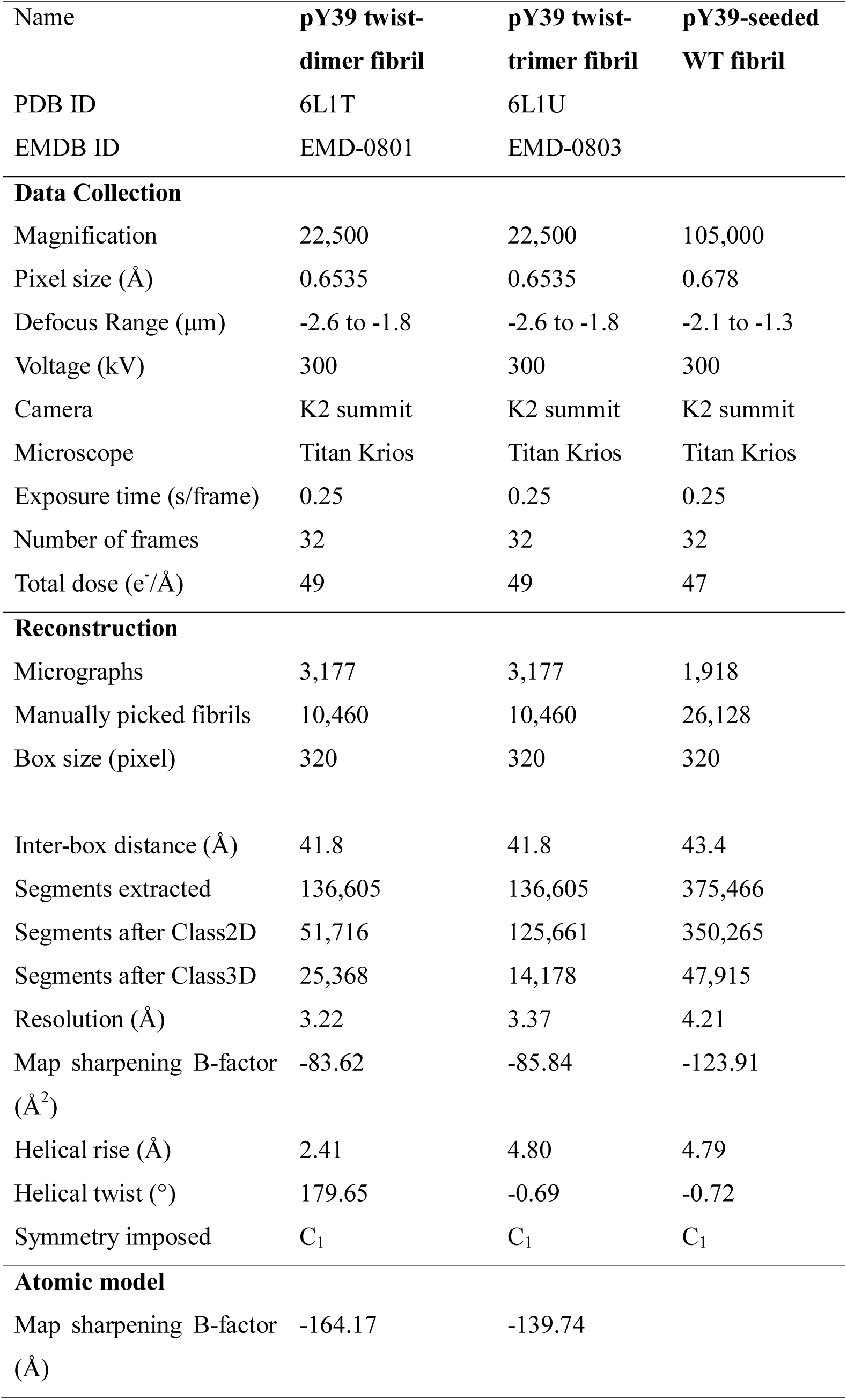
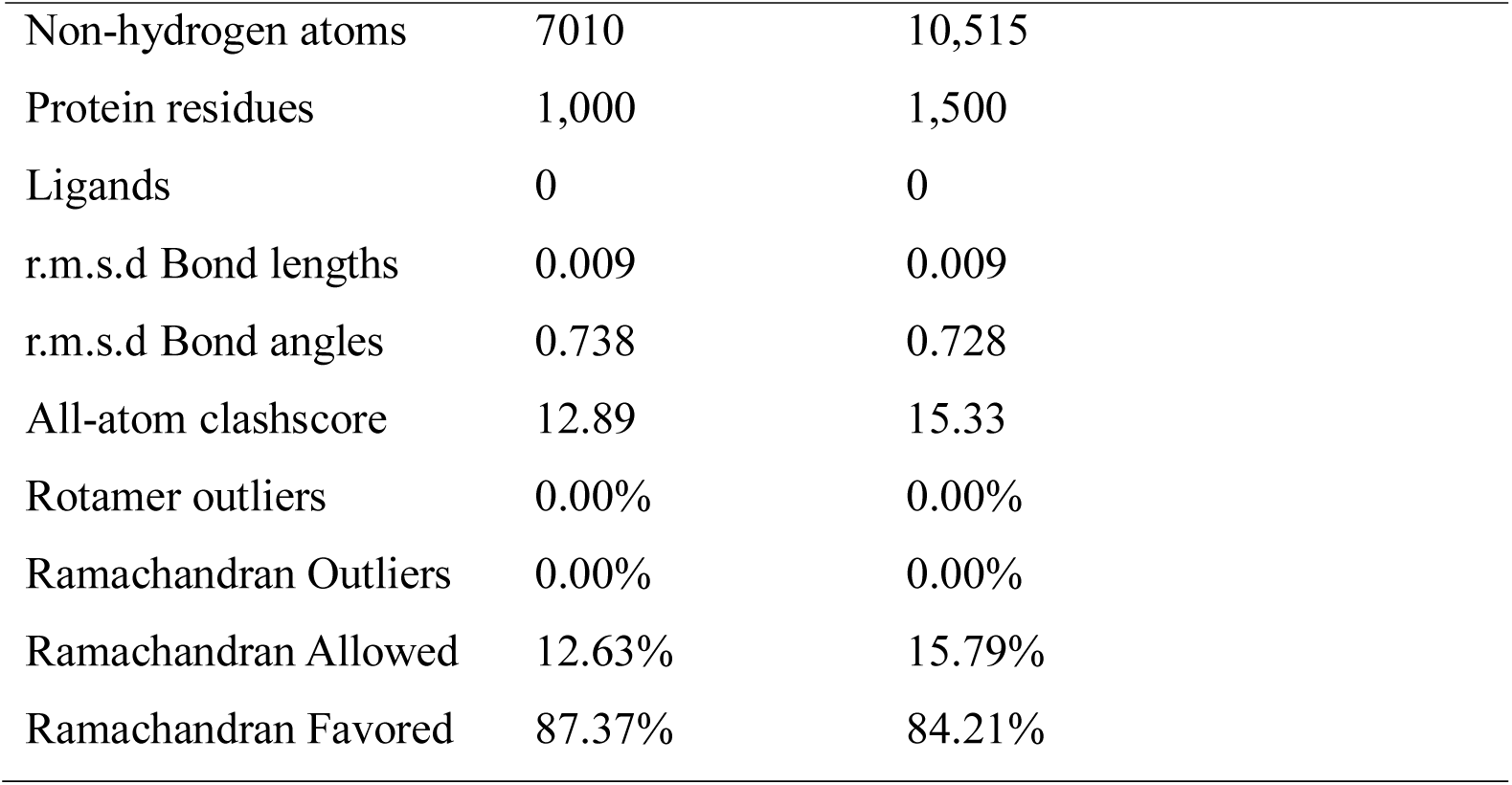
Statistics of cryo-EM data collection and refinement.

| Name | pY39 twist-dimer fibril | pY39 twist-trimer fibril | pY39-seeded WT fibril |
| --- | --- | --- | --- |
| PDB ID | 6L1T | 6L1U |  |
| EMDB ID | EMD-0801 | EMD-0803 |  |
| **Data Collection** |  |  |  |
| Magnification | 22,500 | 22,500 | 105,000 |
| Pixel size (Å) | 0.6535 | 0.6535 | 0.678 |
| Defocus Range (µm) | -2.6 to -1.8 | -2.6 to -1.8 | -2.1 to -1.3 |
| Voltage (kV) | 300 | 300 | 300 |
| Camera | K2 summit | K2 summit | K2 summit |
| Microscope | Titan Krios | Titan Krios | Titan Krios |
| Exposure time (s/frame) | 0.25 | 0.25 | 0.25 |
| Number of frames | 32 | 32 | 32 |
| Total dose (e<sup>-</sup>/Å) | 49 | 49 | 47 |
| **Reconstruction** |  |  |  |
| Micrographs | 3,177 | 3,177 | 1,918 |
| Manually picked fibrils | 10,460 | 10,460 | 26,128 |
| Box size (pixel) | 320 | 320 | 320 |
| Inter-box distance (Å) | 41.8 | 41.8 | 43.4 |
| Segments extracted | 136,605 | 136,605 | 375,466 |
| Segments after Class2D | 51,716 | 125,661 | 350,265 |
| Segments after Class3D | 25,368 | 14,178 | 47,915 |
| Resolution (Å) | 3.22 | 3.37 | 4.21 |
| Map sharpening B-factor (Å<sup>2</sup>) | -83.62 | -85.84 | -123.91 |
| Helical rise (Å) | 2.41 | 4.80 | 4.79 |
| Helical twist (°) | 179.65 | -0.69 | -0.72 |
| Symmetry imposed | C<sub>1</sub> | C<sub>1</sub> | C<sub>1</sub> |
| **Atomic model** |  |  |  |
| Map sharpening B-factor (Å) | -164.17 | -139.74 |  |

|  |  |  |
| --- | --- | --- |
| Non-hydrogen atoms | 7010 | 10,515 |
| Protein residues | 1,000 | 1,500 |
| Ligands | 0 | 0 |
| r.m.s.d Bond lengths | 0.009 | 0.009 |
| r.m.s.d Bond angles | 0.738 | 0.728 |
| All-atom clashscore | 12.89 | 15.33 |
| Rotamer outliers | 0.00% | 0.00% |
| Ramachandran Outliers | 0.00% | 0.00% |
| Ramachandran Allowed | 12.63% | 15.79% |
| Ramachandran Favored | 87.37% | 84.21% |

**Fig. 2.**
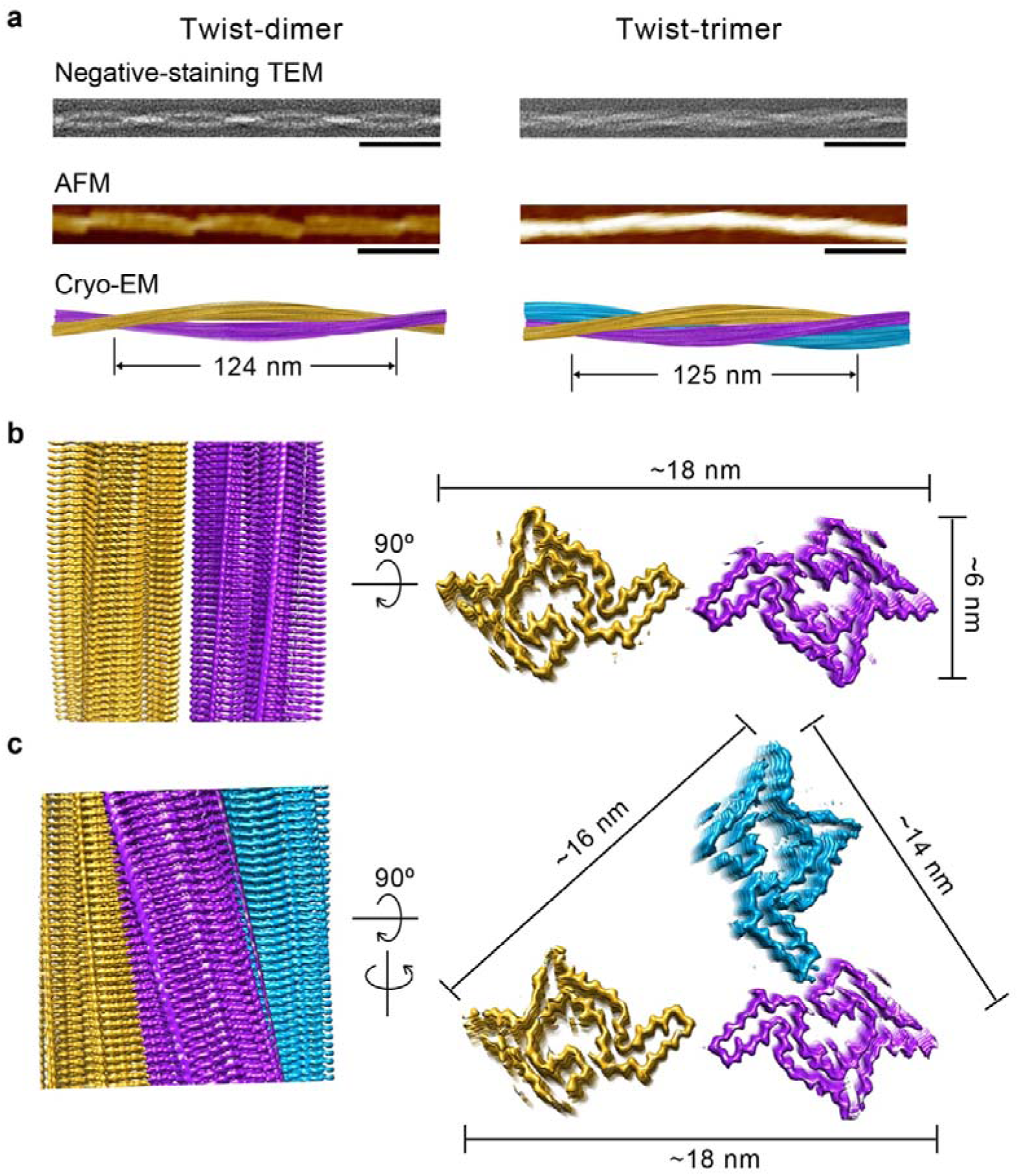
Structure determination of pY39 α-syn fibrils. (*a*) The negative-staining TEM images, AFM images and cryo-EM reconstruction density maps of twist-dimer and twist-trimer fibrils. Scale bar = 100 nm. Half pitches of the fibrils are indicated. (*b*, *c*) Cryo-EM density maps of the pY39 α-syn twist-dimer fibril (*b*) and twist-trimer fibril (*c*). Fibril widths are indicated. Protofilaments are colored individually.

### Relation between the different polymorphs of pY39 α-syn fibrils

According to the density maps, we were able to unambiguously build the atomic model for the two twisted polymorphs (Fig. 3*a*). We aligned the α-syn monomers from protofilament A of the two polymorphs and found that they are nearly identical with an r.m.s.d. of 0.336 Å (93 out of 100 Cα atoms are aligned) (*SI Appendix*, Fig. S10*a*). Further, we aligned the α-syn dimers from protofilaments A and B of the two polymorphs and found that they are also nearly identical with an r.m.s.d. of 0.406 Å (194 out of 200 Cα atoms are aligned) (*SI Appendix*, Fig. S10*b*).

**Fig. 3.**
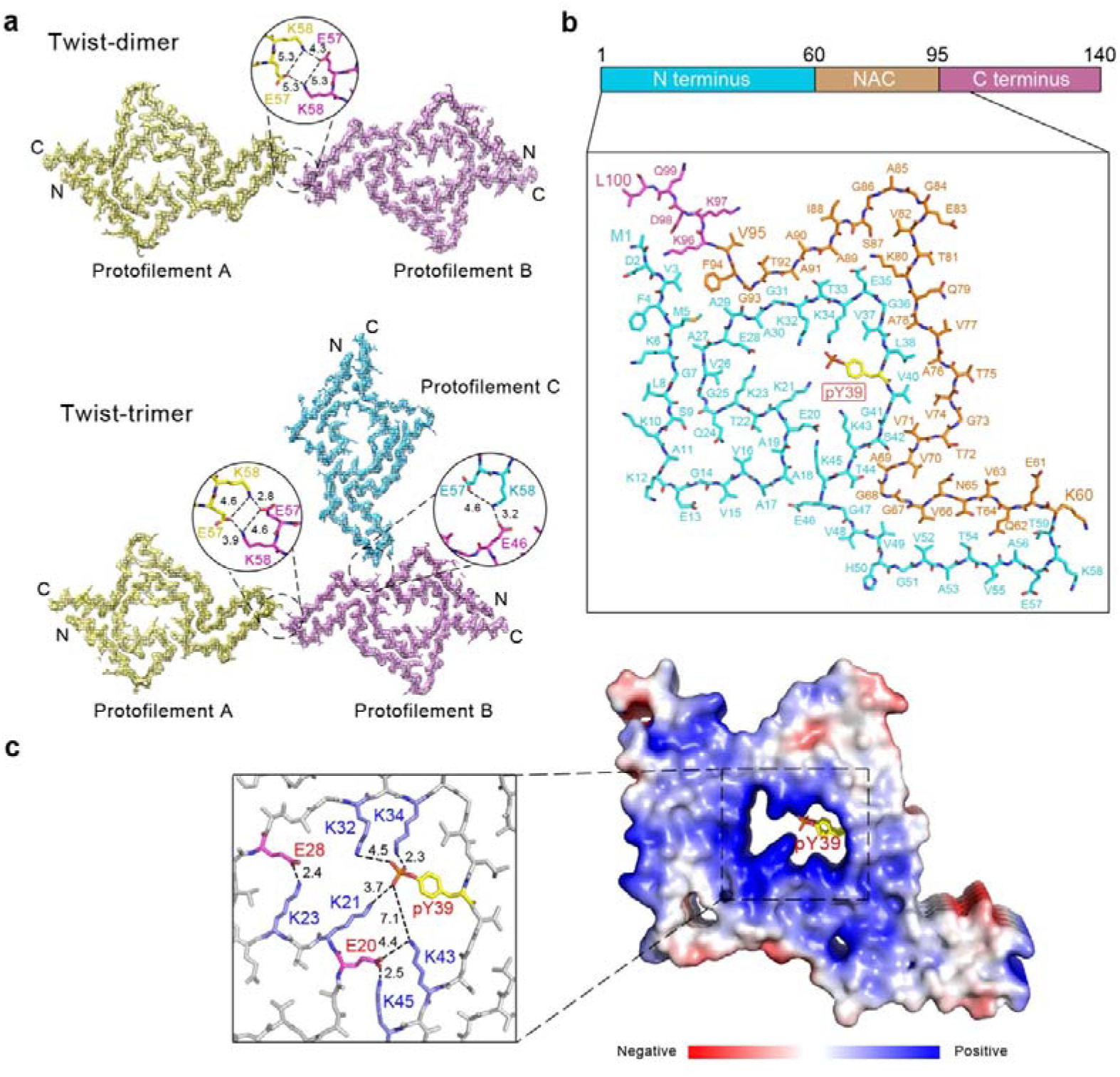
Cryo-EM structures of the pY39 α-syn fibrils. (*a*) Density maps with structural models for twist-dimer (upper) and twist-trimer (lower) fibrils. One dimer/trimer is shown and colored by molecules. Molecular interfaces are shown in the zoom-in views with involved residues and interactions labeled. Distances are shown in Å. (*b*) The structure of pY39 α-syn that is conserved in different twisted polymorphs is shown and colored by domains. The domain organization of α-syn is shown on top. Each individual residue is labeled. (*c*) Electrostatic surface of 5 layers of pY39 α-syn fibril. The central positive hole is shown in the zoom-in view with the charged residues and electrostatic interactions labeled. Glu is in red; Lys is in blue; pY39 is in yellow; the phosphate group of pY39 is in red. Distances are in Å.

The interfaces between protofilaments A and B in twist-dimer and twist-trimer are the same, composed of salt bridges formed by E57 and K58 (Fig. 3*a*). If not considering the pseudo 2-fold symmetry, the helical twist of the twist-dimer fibril would be −0.70° and the helical rise would be 4.81 Å, which are nearly identical to the parameters of the twist-trimer fibril (−0.69° and 4.80 Å) (Table 1). These results indicate that the twist-trimer fibril is formed by an additional protofilament (protofilament C) twining around the twist-dimer fibril. Indeed, we observed a successive appearance of twist-dimer and twist-trimer during the fibril growth by TEM (*SI Appendix*, Fig. S11*a*). Moreover, we observed the coexistence of twist-dimer, twist-trimer and straight polymorphs in a single fibril (*SI Appendix*, Fig. S11*b*). In addition, the width of the protofilament of straight fibril is ~6.1 nm measured from the 2D class averages (*SI Appendix*, Fig. S8), which is very close to the width of the conserved protofilament of twisted fibrils. These results indicate that the polymorphism of pY39 α-syn fibril may derive from different packing of a conserved protofilament in the process of fibril maturation.

### pY39 drives the formation of a distinctively large fibril core

In the cryo-EM fibril structure, pY39 α-syn folds into a hook-like architecture composed of resides 1-100 with the entire N-terminus, NAC domain and a short segment of the C-terminus involved (Fig. 3*b*), which presents the largest fibril core of α-syn reported so far. Impressively, in the center of the hook head, lysine residues (K21, K23, K32, K34, K43 and K45) of the N-terminus form a positive hole, which is neutralized by E20, E28 and the phosphate group of the pY39 (Fig. 3*c*). A network of electrostatic interactions is formed between E28/K23, E20/K43/K45 and pY39/K21/K32/K34 (Fig. 3*c*). Residues V52-V66 form a hetero-steric zipper and reach out to mediate inter-protofilamental interactions for the further assembly of mature fibrils (Fig. 3*a* and 3*b*).

The overall topology of the pY39 structure is distinct from any known structures of α-syn fibrils formed by full-length α-syn (22–26) (Fig. 4*a*) or C-terminus truncated WT α-syn (27), or mutant α-syn (28–31), except that the out-reaching steric zipper of pY39 fibril also exists in the structure of polymorph 2a WT fibril reported lately (26) (Fig. 4*b*). Different polymorphs of the α-syn fibril cores reported previously are composed by a common segment covering residues approximately from 37-99 (polymorph 1b is even smaller). This segment is able to fold into various compact structures for the polymorphic formation of α-syn amyloid fibrils (Fig. 4*a*). In contrast, this segment is relatively stretched in the fold of pY39 fibril, and together with the rest of the N-terminal residue (residues 1-36), forms a distinctly large fibril core (Fig. 4*a*).

**Fig. 4.**
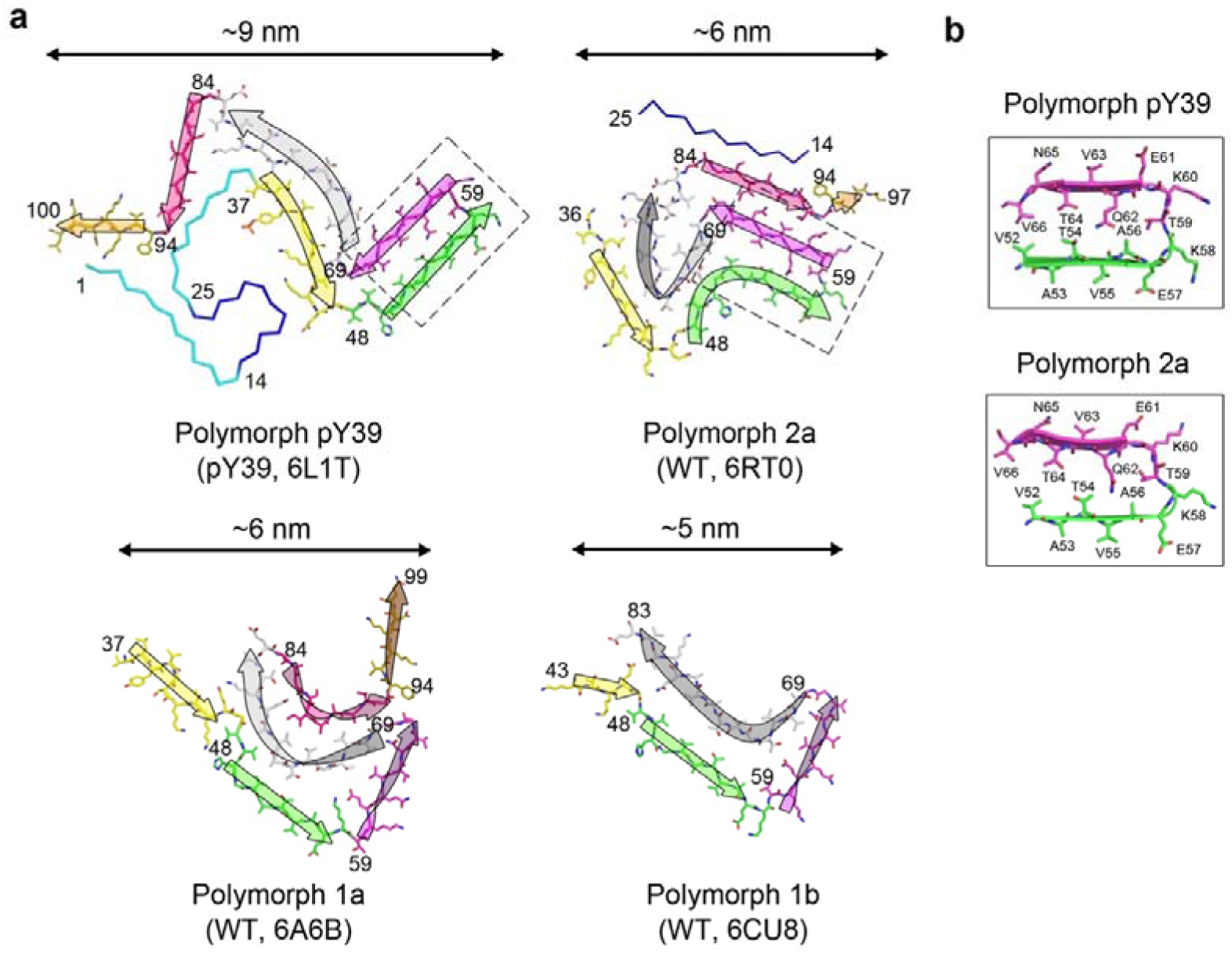
Comparison of polymorphic α-syn fibril structures. (*a*) Topology of α-syn in different polymorphs of amyloid fibrils. α-Syn variants and PDB IDs are shown in the parentheses. The sequence involved in the WT fibril cores is shown in sticks and colored regionally. The N-terminus that is usually flexible in the WT fibril but involved in the core of the pY39 fibril is shown in ribbons and colored in cyan and blue. The arrows indicate the topology of the structures. Flip of the arrow indicates flip of the amino acid chain. Selected residue numbers are shown. The name of WT fibril polymorphs follows Ref 26. Widths of the fibril cores are indicated. Dashed boxes highlight the conserved hetero-steric zipper in pY39 and polymorph 2a. (*b*) A zoom-in view of the conserved steric zipper highlighted in (*a*), shown in cartoon and sticks. Residue numbers are labeled.

### Hydrophilic channel in the center of pY39 α-syn fibril core

The involvement of the entire N-terminus not only enlarges the fibril core, but also significantly increases the proportion of charged residues, including K, D and E (*SI Appendix*, Fig. S12). Note that α-syn does not contain arginine. Besides, phosphorylation further modifies Y39 into a strong negatively-charged residue. With such a high amount of charged residues, pY39 α-syn remarkably folds with eight charged residues (six lysine and two glutamate) plus pY39 to form a hydrophilic channel in the center of the fibril core, which might be accessible to the solvent (Fig. 5*a*). Actually, we observed isolated round density in the channel at the local resolution of 2.9 Å (Fig. 5*b* and *SI Appendix*, Fig. S9*a*), which may represent water molecules hydrogen-bonding with the side chains of pY39, K21 and K43. Thus, the pY39 α-syn fibril may contain an inner and an outer surface to contact with the environment (Fig. 5*c*).

**Fig. 5.**
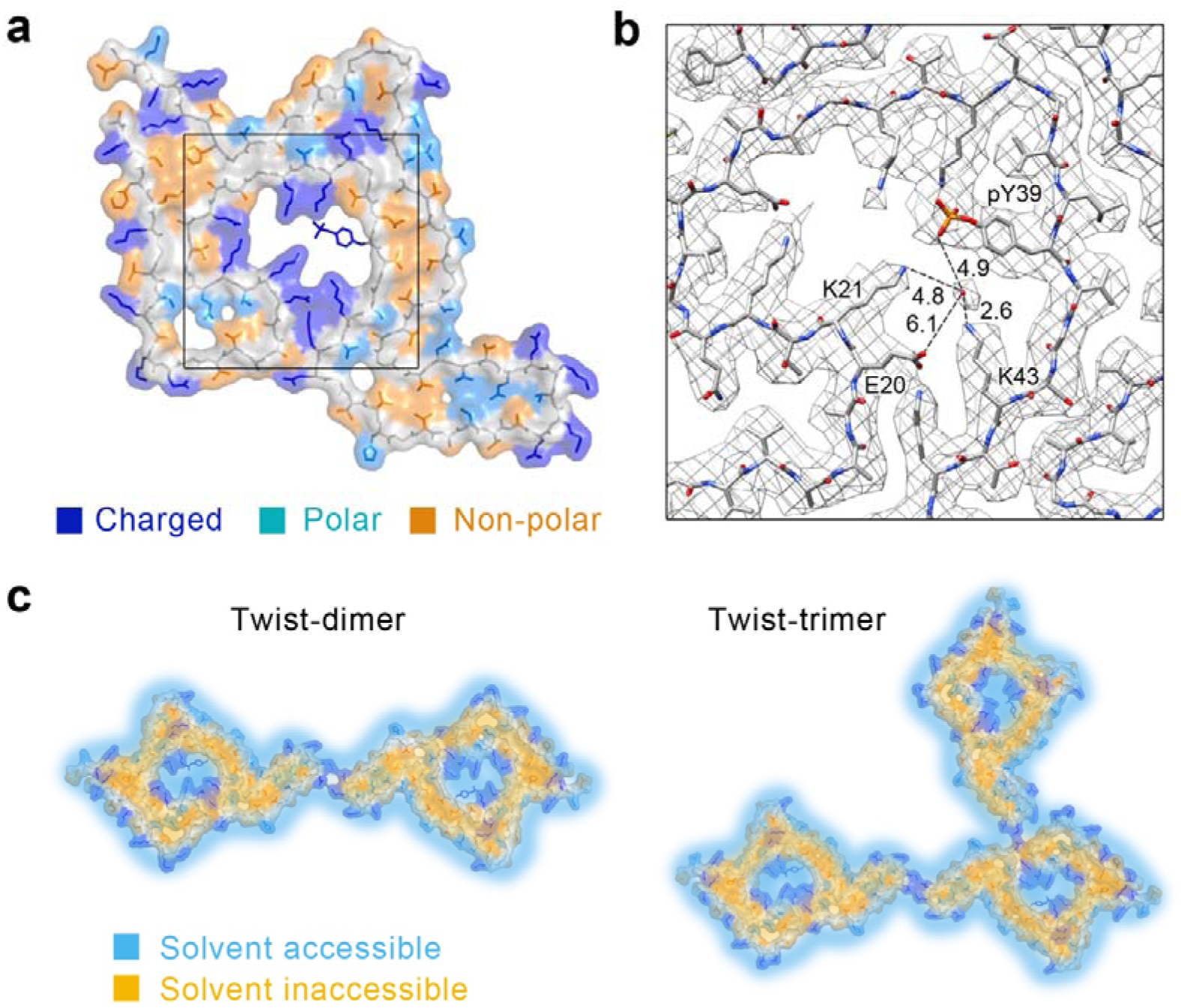
Hydrophilic central channel of pY39 α-syn fibril. (*a*) The surface of the pY39 α-syn is shown. The side chains of charged residues are colored in blue; polar residues are in light blue; non-polar residues are in orange. The main chain is colored in gray. Density map of the framed region is shown in (*b*). (*b*) Density map of the central hole shows the density of water. Distances are in Å. (*c*) The potential solvent-accessible (light blue) and inaccessible (orange) surfaces of the twist-dimer and twist trimer fibrils are shown.

### pY39 α-syn fibrils are more resistant to protease digestion

We used trypsin to digest the WT and pY39 α-syn fibrils, respectively. Of note, since trypsin cleaves proteins at K or R, accordingly to the pY39 structure, only one cleavage site (K102) is outside of the fibril core and most available for trypsin digestion. Indeed, upon trypsin digestion, the pY39 fibril generated only one major band of cleaving product, which covers residues 1-102 analyzed by LC-MS/MS (*SI Appendix*, Fig. S13*a*). In contrast, the WT fibril was digested into several fragments. Slower degradation of pY39 fibril was also observed by protease K (PK) digestion (*SI Appendix*, Fig. 13*b*). In addition, pY39 and WT α-syn monomers were used to confirm the activity of the proteases (*SI Appendix*, Fig. S13*c*). These results indicate that phosphorylation at Y39 may cause α-syn fibrils to be more difficult for clearance in cells.

### pY39 PFFs cross-seed WT α-syn to form amyloid fibrils featuring the WT fibril structure

To understand the potential for the spread of pY39 fibrils, we sought to see if the pY39 fibrils can cross-seed the fibril formation of WT α-syn. We monitored the aggregation kinetics of WT α-syn in the presence of pY39 PFFs by the ThT assay. Intriguingly, the result showed that similar to WT PFFs, pY39 PFFs effectively seeded the amyloid aggregation of WT α-syn (Fig. 1*b* and 6*a*). However, it is difficult to imagine that pY39 PFFs could possibly template non-phosphorylated α-syn into the same fibril structure. Although we did not find WT fibril morphologies in the pY39 fibrils during cryo-EM data processing, there is still a possibility that a trace amount of the WT fibril structures were formed by pY39 α-syn, which might carry out the seeding effect. To rule out this possibility, we seeded WT α-syn aggregation with a gradient of low percentages of WT PFFs. The ThT result showed that the seeding effect decreased as the PFF concentration decreased; in particular, <0.05% WT PFFs, which is equivalent to <5% WT fibril structure in 1% pY39 PFFs, showed apparently weaker seeding effect than that of 1% pY39 PFFs (Fig. 6*a* and *SI Appendix*, Fig. S14). Moreover, there is certainly less than 5%, if not none, WT fibril structure in the pY39 fibrils based on the cryo-EM data (*SI Appendix*, Fig. S3*e*). Thus, these results demonstrate that pY39 fibrils can indeed cross-seed the fibril formation of WT α-syn.

**Fig. 6.**
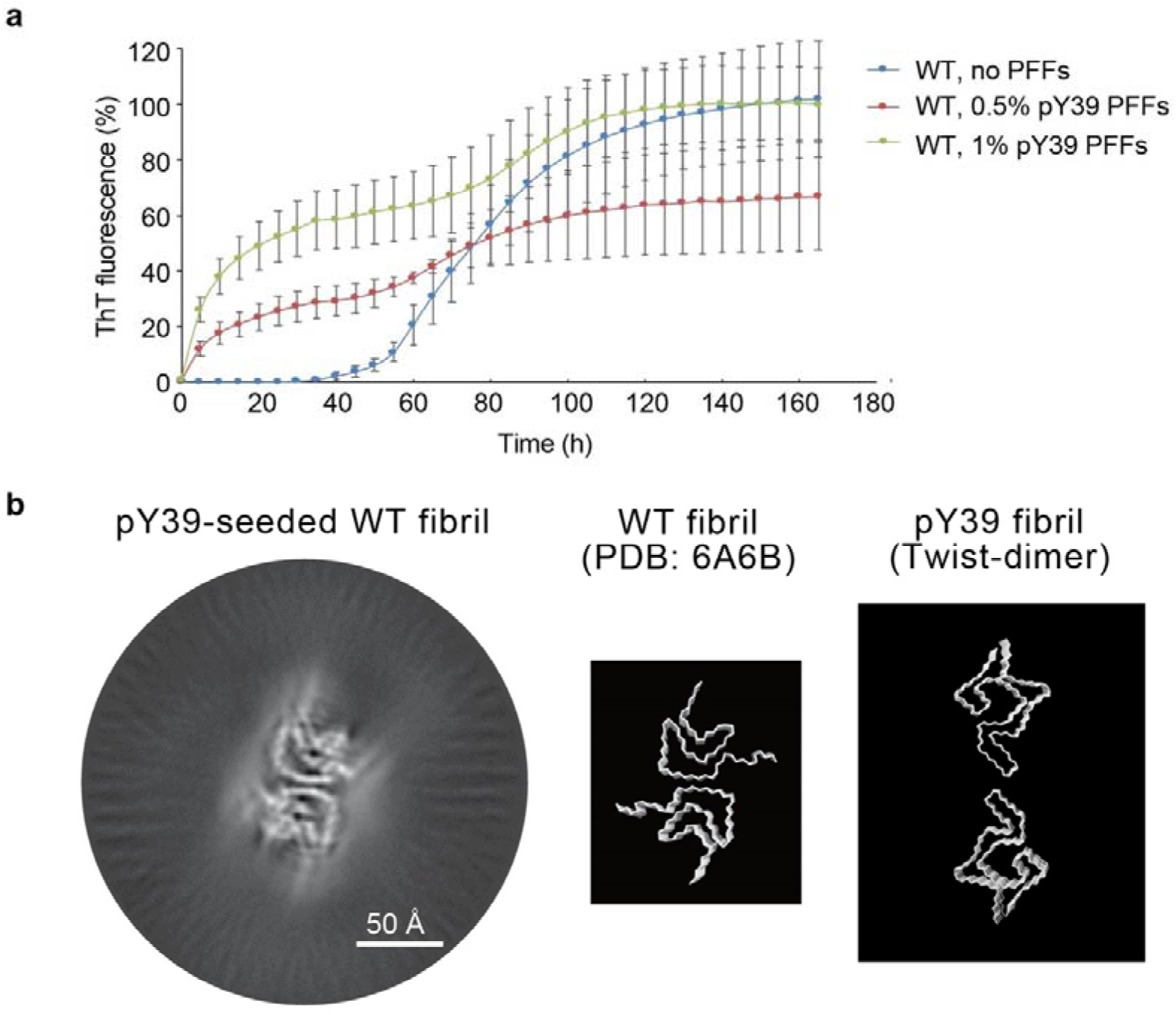
pY39 PFFs cross-seed WT α-syn fibril formation. (*a*) ThT kinetic assay of the pY39-seeded WT α-syn aggregation. Mol% of added PFFs are indicated. Data shown are mean ± s.d., n=3. (*b*) Cryo-EM 3D reconstruction of the pY39-seeded WT α-syn fibril (left). Structural models of WT α-syn fibril (PDB ID: 6A6B) and pY39 twist-dimer are shown on the right.

To understand whether and how WT α-syn lacking the phosphate group at Y39 could possibly fold into the structure as that in the pY39 fibril, we performed cryo-EM to determine the structure of pY39-seeded WT α-syn fibril. Although the image processing did not allow sufficiently high resolution for model building, the result of 3D reconstruction showed that the cross-seeded fibril features the structure of WT fibril (polymorph 1a) rather than pY39 fibril (Fig. 6*b*). This result confirms the key role of Y39 phosphorylation in the formation of the pY39 fibril. Without pY39, WT α-syn can hardly form the structure as that in the pY39 fibril. Although the mechanism of pY39 cross-seeding WT α-syn fibril formation remains unknown, the phenomenon is not surprising since cross-seeding effect has been widely found between different amyloid proteins such as between α-syn and Aβ (32), between Aβ and IAPP (33,34), and between Aβ and Tau (35).

## Discussion

In this work, we reveal that Y39 phosphorylation leads to the formation of new polymorphs of α-syn amyloid fibril, which feature a distinctly large fibril core. Structural polymorphism is a common feature of amyloid fibrils formed by different peptides/proteins, such as α-syn, Tau and Aβ (22, 24, 36–40). Different fibril polymorphs may exhibit different pathologies under disease conditions (6, 41). For instance, α-syn fibrils derived from PD and MSA patients exhibit distinctive morphologies and pathologies (6). Thus, structural understanding on different fibril polymorphs is important for the understanding of the complicated pathology of the disease. In recent years, technological advance in cryo-EM enables the structure determination of various full-length amyloid fibrils at the near-atomic resolution (42). Polymorphic core structures have been determined in the WT and mutant α-syn fibrils (22–31), which are commonly composed of segment ~40-100 or smaller. Both the C-terminus and the majority of the N-terminus remain flexible, and thus are invisible in the cryo-EM structures. The C-terminus is highly enriched in acidic residues, and the N-terminus is enriched in alkaline residues. The high proportion of charged residues precludes the engagement of the two termini in the fibril core structure. Intriguingly, a recent structural study on human brain-derived α-syn fibril showed that *in vivo* cofactors can participate into α-syn fibril structure and introduce more N-terminal region into the fibril core (43), which recalls the observation in the pY39 fibril. Phosphorylation at Y39 involves the entire N-terminus into the fibril core and rearranges the fibril core structure. pY39 changes the electrostatic property of the N-terminus and stabilizes multiple lysine residues to form a hydrophilic channel in the center of the fibril core. Thus, the patient-derived and pY39 α-syn fibril structures together suggest the important role of the N-terminus in the pathological fibril formation of α-syn. In addition, pY39 fibril structure highlights the importance of PTM in defining the structure and pathology of amyloid fibrils.

Phosphorylation widely exist in different amyloid proteins such as α-syn, Tau, Aβ and FUS (10, 13, 44, 45). However, whether and how phosphorylation influences the fibril formation is poorly understood. Recently, a fibril structure of phosphorylated Ser8 (pS8) Aβ40 has been determined by solid-state NMR, while the segment containing pS8 is flexible due to limited NMR constraints (45). A recent cryo-EM study on α-syn reconstructed a low-resolution density map for the pS129 α-syn fibril, which at that resolution is similar to that of the WT fibril (24). Tau is hyperphosphorylated at multiple sites (e.g. S202, S262, S356 and S396) in the progress of amyloid aggregation (10). Whereas, phosphorylation cannot be determined in the cryo-EM structures of Tau fibrils derived from different tauopathies, although the fibrils are indeed phosphorylated probed by antibodies (36, 37, 40). A possible explanation is that phosphorylation of native fibrils is heterogeneous, and thus invisible in cryo-EM structures. As well, the phosphorylation sites that are not involved in the fibril core, are unable to be determined by cryo-EM. In this work, we use semi-synthetic α-syn with site-specific phosphorylation at Y39 to form amyloid fibrils *in vitro*. The pathology and protease resistance of the fibril are consistent to that of native pY39 α-syn aggregates (16, 17), which indicates the biological relevance of the synthetic fibril. Moreover, the unique structure of pY39 α-syn fibril, in particular the key role of pY39 in stabilizing the fibril structure via electrostatic interactions may limit the possibility of alternative structures. Indeed, although pY39 PFFs can seed WT fibril formation, the seeded fibrils still feature WT rather than pY39 fibril structure. Thus, although prepared *in vitro*, the unique pY39 fibril structure presented in this work is informative for the structural understanding of native pY39 fibril.

Previous studies suggest that pY39 phosphorylation can enhance α-syn pathology and protect α-syn against degradation via the autophagy and proteasome pathways in mouse cortical neurons (16). Consistently, we found that the pY39 fibril exhibits enhanced propagation and cytotoxicity in the neuronal model and tends to be more resistant to protease cleavage than the WT fibril. The structures of pY39 fibrils may provide a structural explanation for these phenomena. The involvement of the entire N-terminus into the fibril core enlarges the folded structure of α-syn, which may protect α-syn from protease digestion; on the other hand, chaperones that recognize the N-terminus of α-syn (46) may be inefficient to prevent α-syn aggregation or fail to mediate its clearance. Besides, other factors such as the internalization efficiency may also influence the pathology of pY39 fibrils. The enhanced pathology of pY39 fibrils may play a role in the progression of PD and α-synucleinopathies.

## Materials and methods

### Semi-synthesis of pY39 α-syn

To obtain the full-length pY39 α-syn, we performed N-to-C sequential native chemical ligation using three segments including segment A (α-Syn 1-29NHNH_2_), segment B (pY39 α-syn A30C-52NHNH_2_) and segment C (α-Syn A53C-140) (*SI Appendix*, Fig. S1) (19, 47, 48). The segments A and B were manually synthesized by Fmoc-based solid phase peptide synthesis (Fmoc-SPPS). The sequence of segment A is MDVFMKGLSKAKEGVVAAAEKTKQGVAEA-NHNH_2_. The sequence of segment B is CGKTKEGVL[pY]VGSKTKEGVVHGV-NHNH_2_. The segment A and segment B were purified with RP-HPLC and characterized by analytical RP-HPLC and ESI-MS (*SI Appendix*, Fig. S2).

To obtain segment C (α-Syn A53C-140), we fused the SUMO-tag to the N-terminus of α-syn 54-140 and overexpressed the SUMO-fused α-syn 54-140 in BL21 (DE3) *E. coli* cells (48). Protein expression was induced by addition of 1 mM isopropyl-β-D-1-thiogalactopyranoside (IPTG) and incubation of cells for 20 h at 16 °C. Then, the cells were harvested in buffer (50 mM NaH_2_PO_4_, 500 mM NaCl, pH 7.4), and lysed by ultrasonication (twice on ice, each for 20 min). The fusion protein was purified by Ni-NTA. To remove the SUMO tag, the SUMO-C-α-syn 54-140 was treated with protease ULP1 in cleavage buffer (50 mM NaH_2_PO_4_, 500 mM NaCl, pH 8.0) at 16℃ overnight. Then, the SUMO tag and ULP1 were removed using Ni-NTA. The segment C was purified by preparative RP-HPLC with Proteonavi column. Characterization of segment C was achieved by using analytical HPLC and ESI-MS (*SI Appendix*, Fig. S2).

Then, the three segments were N-to-C sequentially linked by native chemical ligation (NCL) for producing the full-length pY39 α-syn. Firstly, segment A and segment B were ligated to produce segment AB (pY39 α-syn 1-52NHNH_2_, A30C). 2 mM Segment A was dissolved in 400 μl 0.2 M sodium phosphate buffer containing 6 M GuHCl (PS buffer) at pH 3.0 and put into ice-salt bath to pre-cool. In order to oxidize the peptide hydrazide to the corresponding azide, 0.05 M NaNO_2_ was added in the reaction for 15 min. To convert the peptide azide to the thioester, 0.2 M MPAA in 400 μl PS buffer was added and then the pH was adjusted to 6.8. To complete ligation reaction, 1.2 eq. segment B was added and reacted at 16 ^o^C overnight. The segment AB was further purified with RP-HPLC and characterized by analytical RP-HPLC and ESI-MS (*SI Appendix*, Fig. S2). Secondly, we ligated segment AB and segment C to obtain segment ABC (pY39 α-syn 1-140, A30C, A53C). The ligation of segments AB and C was similar to that of A and B. The segment ABC was further purified with RP-HPLC and characterized by analytical RP-HPLC and ESI-MS (*SI Appendix*, Fig. S2). Finally, we changed Cys to Ala in segment ABC by chemical desulfurization to obtain the full-length pY39 α-syn protein. Under free radical conditions (0.4 M TCEP, 0.5 mM 2,2′-Azobis[2-(2-imidazolin-2-yl)propane]dihydrochloride (VA-044), and 10% (v/v) 2-methylpropane-2-thiol in 0.2 M PS buffer pH 7.0), the ligation product was desulfurized at 37 °C overnight. pY39 was synthesized in milligram amounts with high purity as assessed by RP-HPLC and ESI-MS (*SI Appendix*, Fig. S2). Note that the N-terminus of the synthesized pY39 α-syn is not acetylated.

### Fibril preparation of WT and pY39 α-syn

Fibrillation was conducted by diluting recombinant non-modified WT α-syn and synthetic pY39 α-syn to 100 μM in buffer (50 mM Tris, pH 7.5, 150 mM KCl, 0.05% NaN_3_). WT and pY39 α-syn are incubated at 37 °C with constant agitation at 900 rpm in ThermoMixer (Eppendorf) for a week, respectively. The fibrils were sonicated for 30 s (1s on, 1s off) on ice at 20% power by JY92-IIN sonicator (Xinyi Sonication Equipment Company, Ningbo, China). The sonicated fibrils (0.5%, v/v) were mixed with 100 μM α-syn monomer, and further incubated at 37 °C with agitation (900 rpm) for two weeks to obtain fibrils for cryo-EM and AFM experiments. To prepare α-syn PFFs for the primary neuron transmission and protease digestion experiments, the fibrils used for cryo-EM were concentrated by centrifugation (14,000 rpm, 25 °C, 45 min), washed with PBS for three times, and sonicated again using the same sonication condition.

### ThT kinetic assay

pY39 and WT α-syn PFFs were prepared by sonication at 20% power on ice for 15 times (1 s/time, 1 s interval) by JY92-IIN sonicator (Xinyi Sonication Equipment Company, Ningbo, China). 50 μM of pY39 or WT α-syn monomer in buffer (50 mM Tris, pH 7.5, 150 mM KCl, 0.05% NaN_3_) was incubated with or without PFFs in clear 384-well plates (Thermo Scientific Nunc). Final concentration of 10 μM ThT was added to the reaction mixture. The plate was detected by a FLUOstar Omega Microplate Reader (BMG LABTECH). Fluorescence was measured at 440 nm (excitation) and 485 nm (emission) wave-lengths, with a bottom read. Three replicates were performed for each sample.

### Negative-staining TEM

An aliquot of 5 μl of fibril solution was loaded onto a glow-discharged grid with carbon support film (Beijing Zhongjingkeyi Technology Co., Ltd.). After 45 s, the grid was washed with 5 μl double-distilled water and 5 μl 3% w/v uranyl acetate. Then 5 μl 3% w/v uranyl acetate was used to stain the sample for 45 s, Filter paper was used to remove the excess solution. The samples were imaged using Tecnai T12 microscope (FEI Company) operated at 120 kV.

### Atomic force microscopy (AFM)

Samples were diluted to a final concentration of 50 μM in the fibril growth buffer (50 mM Tris, 150mM KCl, pH 7.5). 8 μl of the sample were deposited on the surface of freshly cleaved mica at room temperature for 3 min and washed with Milli-Q water for AFM imaging. The samples were recorded in air by using Nanoscope V Multimode 8 (Bruker) in ScanAsyst mode using probe SNL-10. Continuous curves were recorded with a frequency of 65 kHz. Images were acquired at 1024×1024 pixels at a line rate of 1.5 Hz. The height of samples was processed on the NanoScope Analysis software (version 1.5).

### Primary neuronal cultures

Primary cortical neurons were prepared from the cortex of embryonic day (E) 16–E18 Sprague Dawley rats (Shanghai SIPPR BK Laboratory Animals Ltd, China) embryos as previously described (20). In brief, dissociated cortical neurons were plated onto coverslips in 24-well plate at a density of 150,000 cells/well, coated with poly-D-lysine. Neurons were then treated with PBS and WT or pY39 α-syn PFFs at 8 days in vitro (DIV) and collected for immunocytochemistry at 14/18/22 days post-treatment. All rat experiments were performed according to the protocols approved by Animal Care Committee of the Interdisciplinary Research Center on Biology and Chemistry (IRCBC), Chinese Academy of Sciences (CAS). Each experiment was repeated for more than three times.

### Immunofluorescence imaging

Primary cortical neurons were fixed with 4% paraformaldehyde and 4% sucrose in PBS, then permeabilized with 0.15% Triton X-100. Fixed coverslips were blocked with 5% bovine serum albumin in PBS for 1 h at room temperature. Neurons were incubated with anti-α-syn (Cell Signaling Technology, #2642, 1:1,000 dilution), anti-p129-α-syn (Abcam, ab51253, 1:1,000 dilution), anti-MAP2 (Abcam, ab5392, 1:1,000 dilution), anti-Ub (Santa Cruz Biotechnology, sc-8017, 1:200), anti-P62 (Santa Cruz Biotechnology, sc-28359, 1:200) antibodies, respectively. After washing with PBS for three times, neurons were stained with Alexa Fluor 488-, Alexa Fluor 568- or Alexa Fluor 633-conjugated secondary antibodies (Life Technologies, A11039, A11036 and A21070, 1:1,000 dilution, respectively) for 1 h at room temperature. After washing with PBS for additional three times, DAPI was stained with ProLong Gold antifade reagent when coverslips were mounted onto glass slides. The fluorescent images were acquired by confocal microscope. The mean gray value of pS129 α-syn signal for primary cortical neurons were quantified by using ImageJ software.

### Lactate dehydrogenase (LDH) assay

Primary neurons were treated with 100 nM WT and pY39 α-syn PFFs or PBS at DIV8. The neurons were harvested 30 days after treatment. The LDH cytotoxicity assay was performed following the manufacturer’s instructions. The absorbance was recorded at 490 nm and 680 nm (background) with BioTek™ Synergy™ 2 Multi-Mode Microplate Readers (Thermo Scientific). The experiment was independently repeated for more than three times. Statistical significance was calculated by one-way ANOVA using GraphPad Prism.

### Cryo-EM imaging and data processing

pY39 α-syn fibril was diluted to 5 μM. An aliquot of 4 μl pY39 α-syn fibril solution was applied to a glow-discharged carbon electron microscope grid (Quantifoil R1.2/1.3, 300 mesh). After 10 s of incubation, the grid was blotted for 6.5 s, and plunge-frozen into liquid ethane using a FEI Vitrobot Mark IV. Cryo-EM data were collected on a FEI Titan Krios microscope operated at 300 kV with magnification of 22,500×. Super-resolution mode was used for a Gatan K2 Summit camera to record the cryo-EM micrographs with a pixel size of 0.6535 Å. Defocus values were set from −2.6 to −1.8 μm. 32 dose-fractionated frames were taken for each micrograph and the electron dose rate was set at ~6.12 e^−^/s/Å^2^ with a total exposure time of 8 s (0.25 s per frame). Hence, the total dose is ~49 e^−^/Å^2^ per micrograph. Data was collected by the Serial EM software.

As for image pre-processing, MotionCorr2 was used to correct beam induced sample motion and implement dose weighting for all 32 frames (49). The pixel size was further binned to 1.307 Å. CTFFIND4.1.8 was used to estimate the defocus values of dose-weighted micrographs (50).

RELION3.0 was used for manual picking, particle extraction, 2D classification, 3D classifications, 3D refinement and post-processing (51). 10,460 filaments were picked manually from 3,177 micrographs. Particles were extracted into 686 and 320 pixel boxes using the 90% overlap scheme. The fibrils of pY39 straight, twist-dimer and twist-trimer were separated by 2D classification.

### Helical reconstruction

AFM measurement, cryo-EM micrographs and 2D classification were used together to calculate an initial twist angle. 4.8 Å was used as an initial value of helical rise. Local searches of symmetry in 3D classification was used to find the final twist angle and rise value. By 2D class averages of the twist-dimer filament segments, a staggered arrangement was observed between the subunits on the two sides of the filament. Therefore, the two protofilaments of twist-dimer were related by a pseudo-2_1_ axis. Accordingly, the rise value of pY39 twist-dimer was re-set to 4.8/2=2.4 Å and the twist angle was re-set to (360° – 0.7°)/2=179.65°.

Several iterations of 2D classification and 3D classification were performed to select particles which belong to the same polymorph. 3D initial model was built by selected 2D classes. 3D classification was performed several times to generate a proper reference map for 3D refinement. 3D refinement of the selected 3D classes with appropriate reference was performed to obtain final reconstruction.

The final map of pY39 twist-dimer was convergence with the rise of 2.41Å and the twist angle of 179.65°. Post-processing was preformed to sharpen the map with a B-factor of −83.6159 Å^2^. Based on the gold-standard FSC = 0.143 criteria, the overall resolution was reported as 3.22 Å. The final map of pY39 twist-trimer was convergence with the rise of 4.80 Å and the twist angle of −0.69°. Post-processing was preformed to sharpen the map with a B-factor of −85.8426 Å^2^. The overall resolution was reported as 3.37 Å.

### Atomic model building and refinement

Phenix.auto_sharpen (52) was used to sharpen the map of both pY39 twist-dimer and twist-trimer. Atomic models were subsequently built into the refined maps with COOT (53). The model of pY39 twist-dimer was built based on the model of WT α-syn fibril (PDBID: 6A6B). To build the pY39 twist-trimer model, a single chain from the twist-dimer structure was extracted and fitted into the density map of twist-trimer. For both twist-dimer and twist-trimer fibrils, a 5-layer model was generated for structure refinement. Structure models were refined by using the real-space refinement program in PHENIX (54).

### Protease digestion of α-syn PFFs

10 μg of α-syn PFFs in PBS (PH 7.4) were incubated for 30 min at 37 ℃ in the presence of trypsin (Promega) or proteinase K (Invitrogen). Digestion reaction was terminated by adding 1 mM PMSF. The samples were boiled with SDS-loading buffer for 15 min, and then resolved on 4%-20% Bis-Tris gels (GenScript). The gels were stained with Coomassie brilliant blue and imaged with Image Lab 3.0 (Bio-Rad).

### In-gel digestion and LC-MS/MS analysis

Target bands were cut from the Coomassie blue-stained SDS-PAGE gel. Proteins in the band were digested by trypsin, and then extracted from the gel following the protocol developed by Matthias Mann’s lab (55). Peptides extracted from the gel were analyzed using an on-line EASY-nL-LC 1000 coupled with an Orbitrap Q-Exactive HF mass spectrometer (Thermo Fisher).

### Accession codes

Density maps of the pY39 α-syn fibrils are available at EMDB with entry codes: EMD-0801 (twist-dimer) and EMD-0803 (twist-trimer). The structure models are deposited in the Protein Data Bank with entry codes: 6L1T (twist-dimer) and 6L1U (twist-trimer). The data that support the findings of this study are available from the corresponding authors upon request.

## Acknowledgements

We thank the Center of Cryo-Electron Microscopy (CCEM) in Zhejiang University School of Medicine for help with data collection. This work was supported by the Major State Basic Research Development Program (2016YFA0501902), the National Natural Science Foundation (NSF) of China (91853113, 81661148047 and 31872716), the National Key R & D Program of China (2018YFA0507600 and 2019YFA0904200), the Science and Technology Commission of Shanghai Municipality (18JC1420500), “Eastern Scholar” project supported by Shanghai Municipal Education Commission, the Shanghai Pujiang Program (18PJ1404300), the Shanghai Municipal Science and Technology Major Project (Grant No. 2019SHZDZX02).

## Author contributions

C.L., Y-M.L. and D.L. designed the project. Y-J.L. and J-J.H. semi-synthesized the pY39 α-syn monomer sample, K.Z., Y-J.L., H.L., Y.S. and X.Z. prepared the WT and pY39 fibril sample and performed the Cryo-EM experiments. K.Z. built and refined the structure model. Z.L., H.L. and C.Z. performed the biochemical and cellular assays. Z.L. performed the AFM experiments. All of the authors are involved in analyzing the data and contributed to manuscript discussion and editing. D.L. and C.L. wrote the manuscript.

