## Supplementary material for "Parkinson’s disease-related phosphorylation at Tyr39 rearranges α-synuclein amyloid fibril structure revealed by cryo-EM"

SI Appendix for

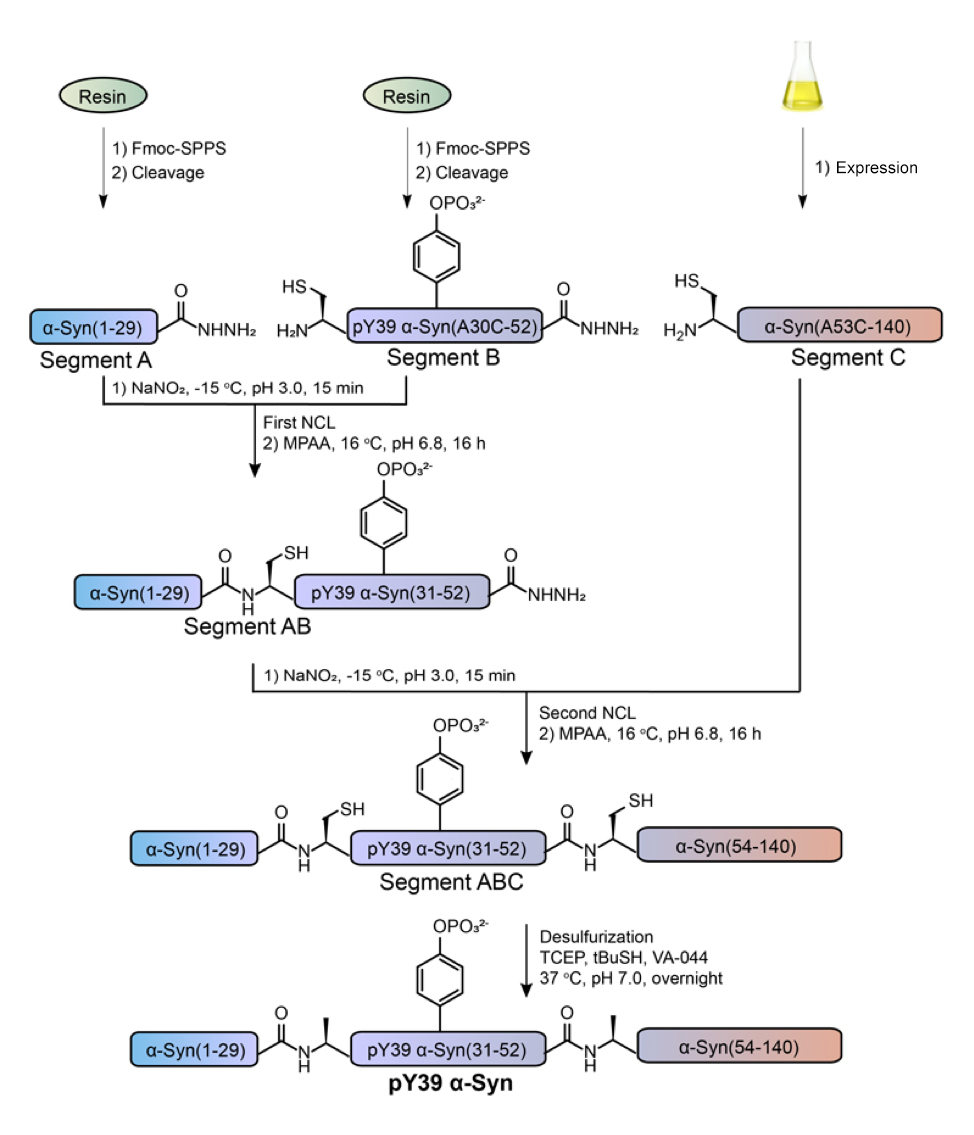

**Figure S1. Workflow of the semi-synthesis of pY39 α-syn.** Reaction conditions are described.

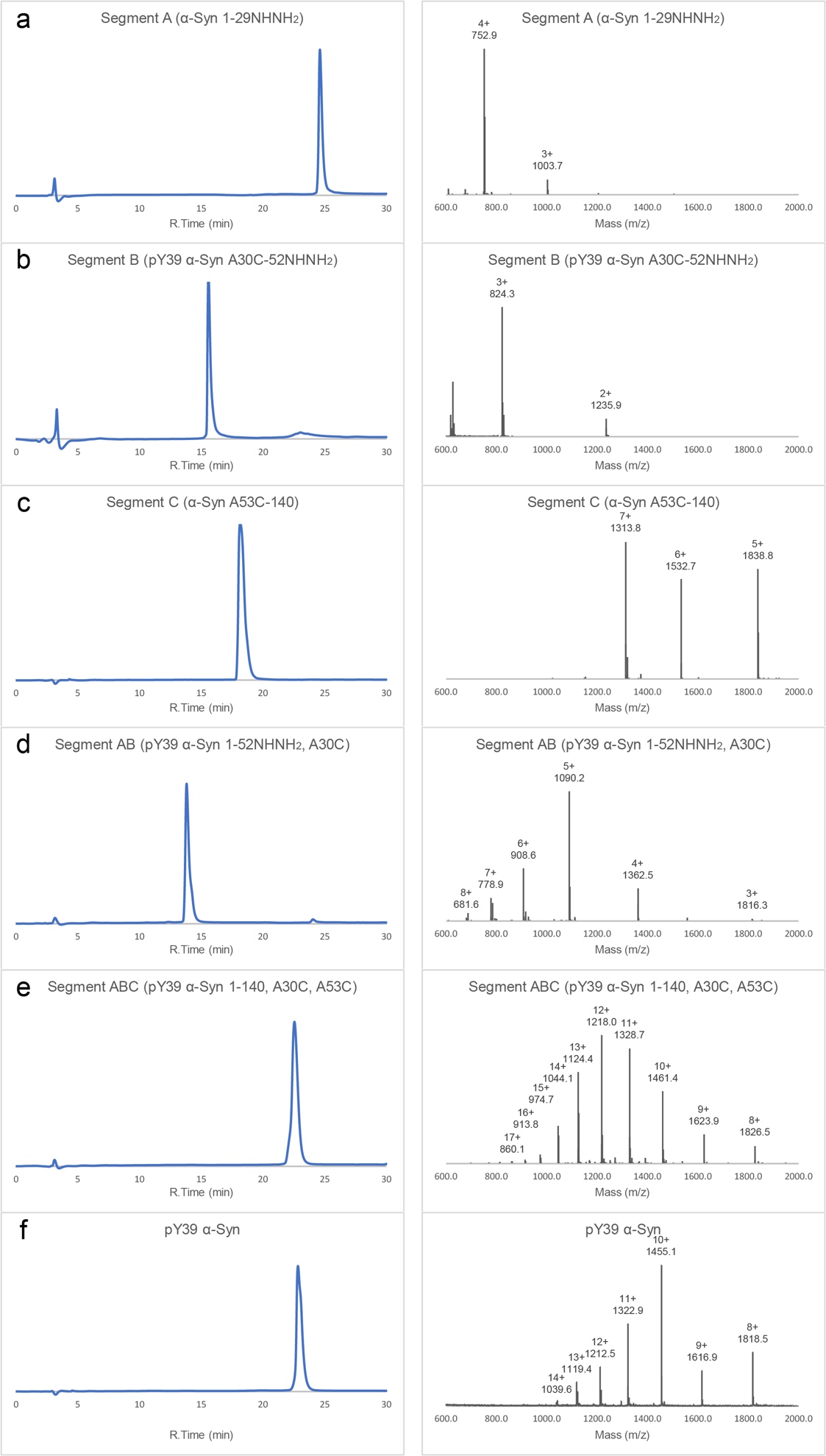

**Figure S2. Characterization of the synthetic pY39 α-syn.** (*a*) Characterization of segment A. RP-HPLC profile (left) of purified segment A showed a peak at 24.62 min. The analysis was run on C18 column with linear gradient of 10-50%B for 30min. ESI-MS (right) of purified segment A (found: 3007.9 Da, calculated: 3008.5 Da). (*b*) Characterization of segment B. RP-HPLC profile (left) of purified segment B showed a peak at 15.57 min. The analysis was run on C18 column with linear gradient of 10-50%B for 30 min. ESI-MS (right) of purified segment B (found: 2469.8 Da, calculated: 2469.8 Da). (*c*) Characterization of segment C. RP-HPLC (left) profile of purified segment C showed a peak at 18.17 min. The analysis was run on Proteonavi column with linear gradient of 30-70%B for 30 min. ESI-MS (right) of purified segment C (found: 9189.6 Da, calculated: 9190.0 Da). (*d*) Characterization of segment AB. RP-HPLC profile (left) of purified segment AB showed a peak at 13.82 min. The analysis was run on Proteonavi column with linear gradient of 30-70%B for 30 min. ESI-MS (right) of purified segment AB (found: 5445.6 Da, calculated: 5446.3 Da). (*e*) Characterization of segment ABC. RP-HPLC profile (left) of purified segment ABC showed a peak at 22.56 min. The analysis was run on Proteonavi column with linear gradient of 30-70%B for 30 min. ESI-MS (right) of purified segment ABC (found: 14604.5 Da, calculated: 14604.3 Da). (*f*) Characterization of pY39 α-syn. RP-HPLC profile (left) of purified pY39 α-syn showed a peak at 22.84 min. The analysis was run on Proteonavi column with linear gradient of 30-70%B for 30 min. ESI-MS (right) of purified pY39 α-syn (found: 14540.4 Da, calculated: 14540.1 Da).

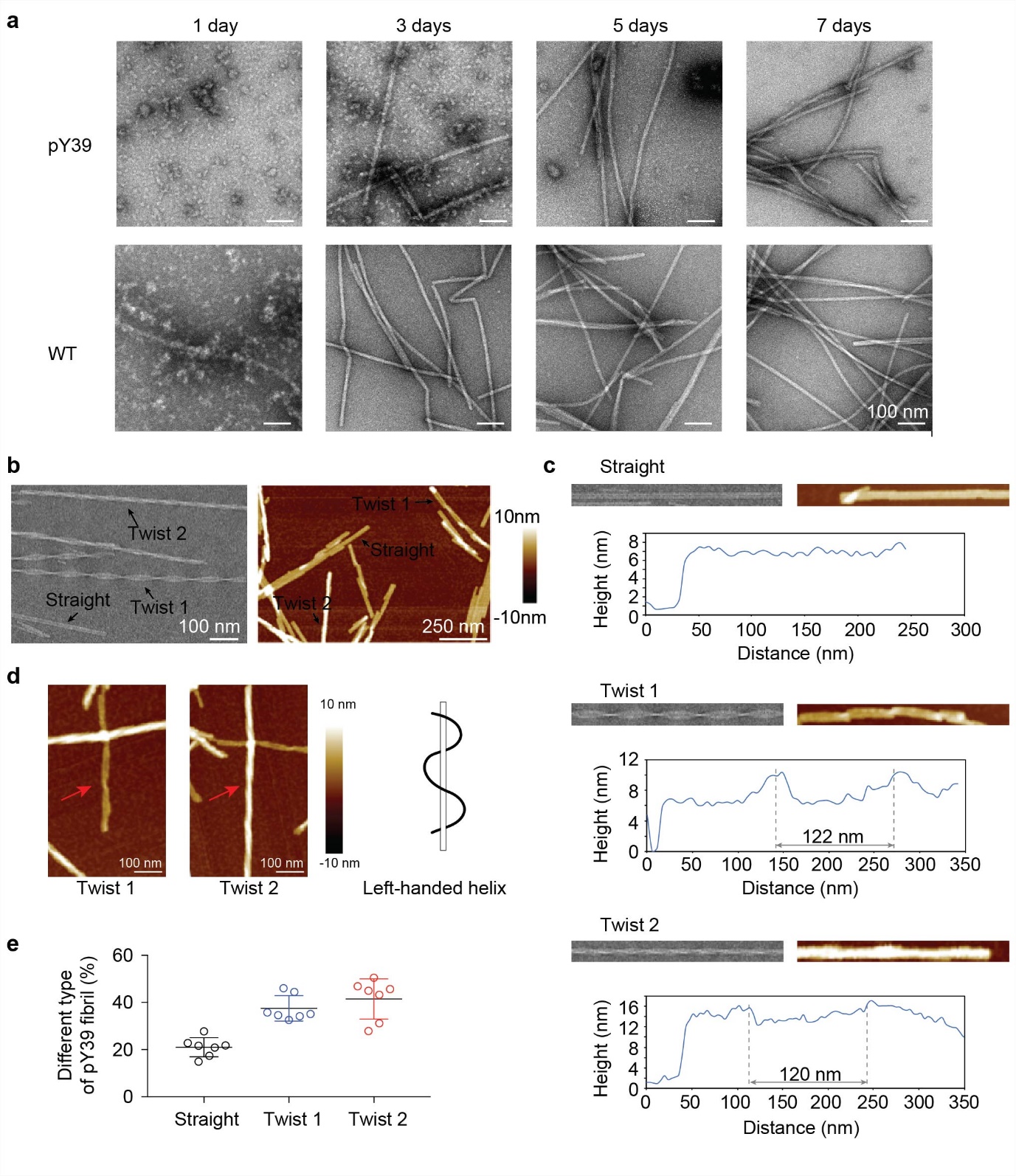

**Figure S3. Polymorphic fibril formation of pY39 α-syn.** (*a*) Time-course tracking of the fibril formation of pY39 and WT α-syn by TEM. (*b*) TEM (left) and AFM (right) images of pY39 fibrils. Three different polymorphs are marked as straight, twist 1 and twist 2. (*c*) Heights of different polymorphic fibrils measured by AFM. Distances between periodic peaks are shown. (*d*) AFM images of the twisted pY39 fibrils. The handedness of the fibrils is indicated. (*e*) The ratio of different polymorphs is calculated from AFM images (n=7).

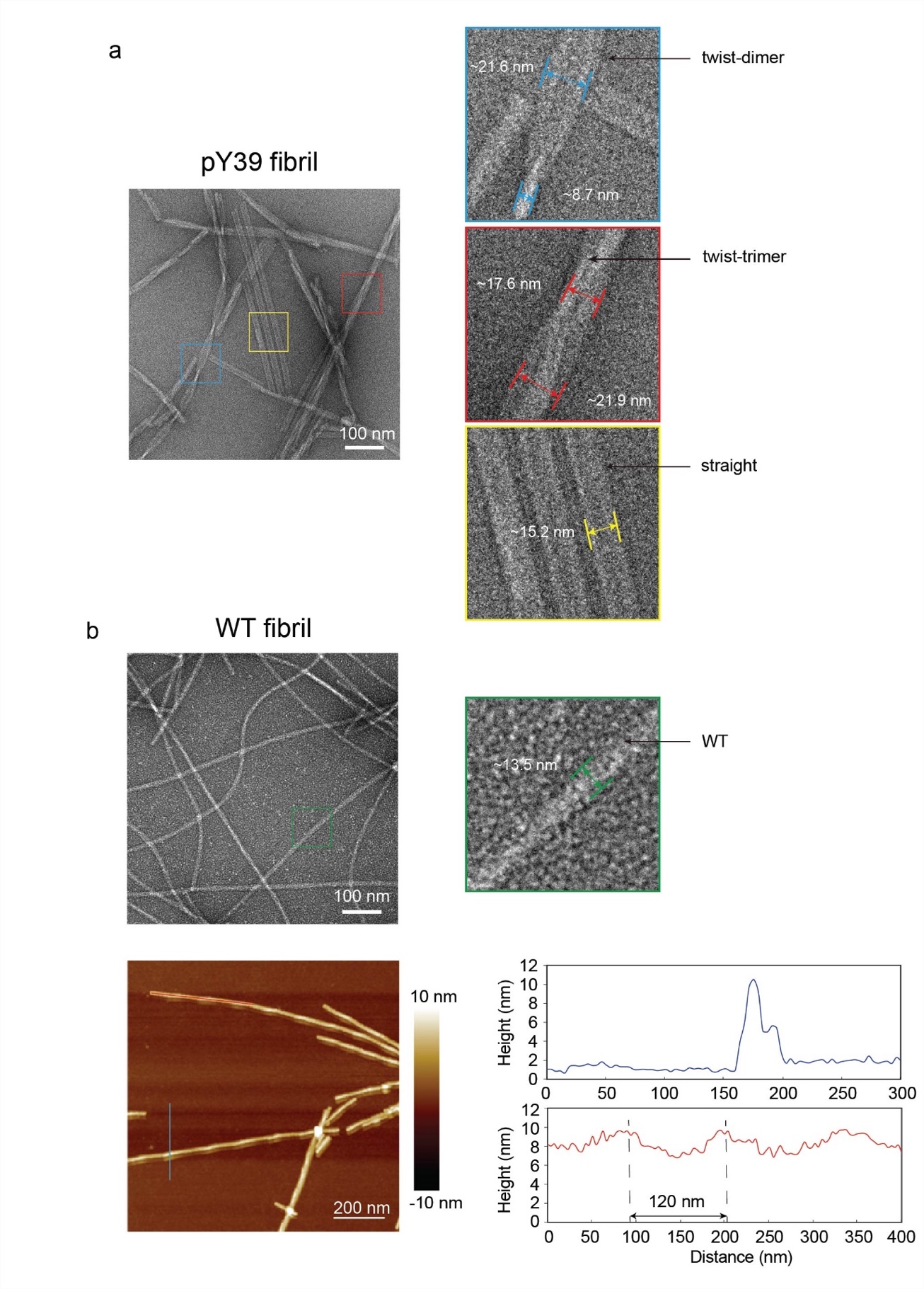

**Figure S4. Distinctive morphologies of pY39 α-syn amyloid fibrils in comparison with the WT fibril formed under the same condition.** (*a*) Negative-staining TEM images of pY39 α-syn fibrils. (*b*) Negative-staining TEM images (top), and AFM image and section analysis of WT fibrils (bottom).

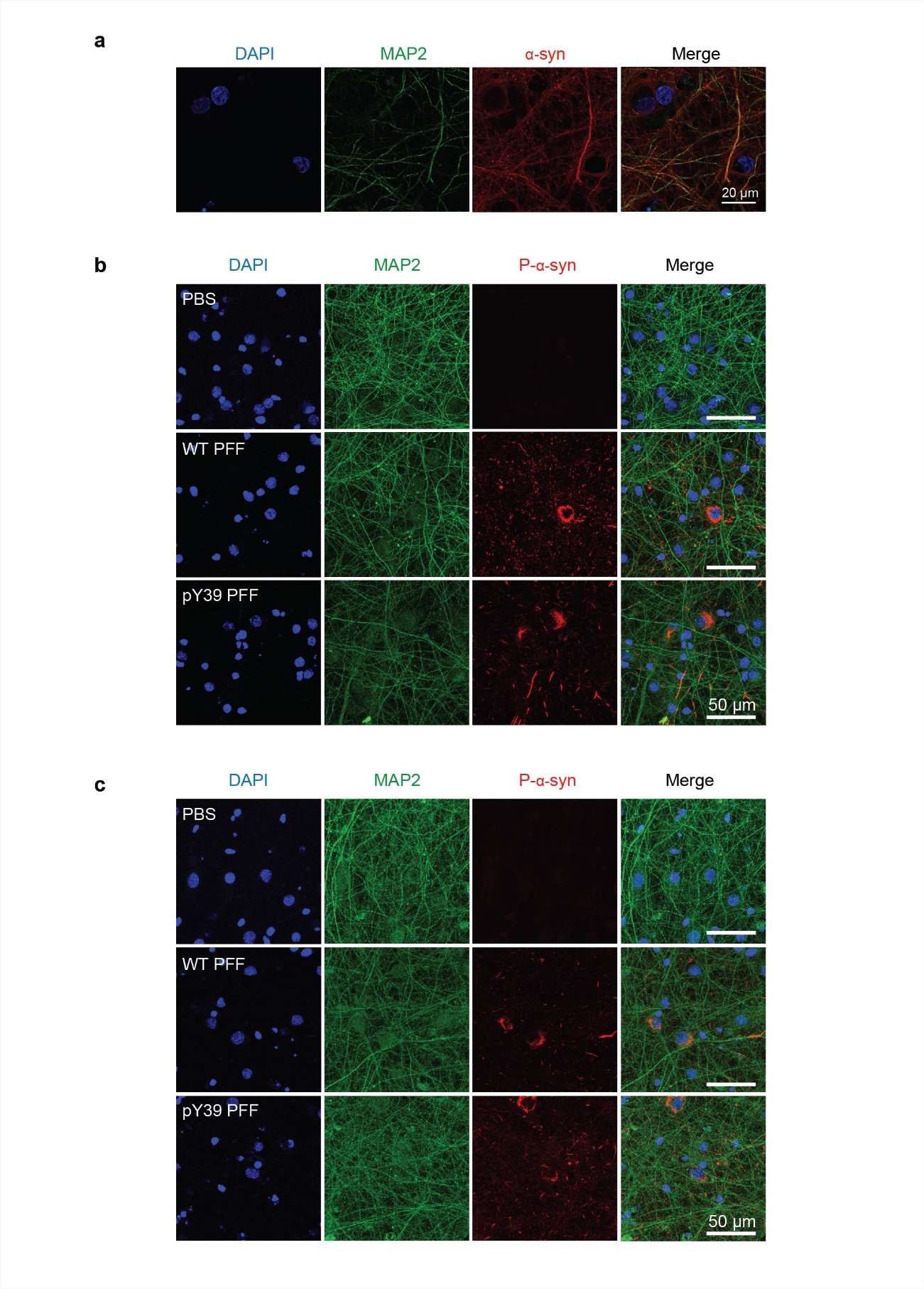

**Figure S5. pY39 α-syn PFFs induce endogenous α-syn aggregation in rat primary cortical neurons.** Confocal microscopic images of fixed neurons stained for DAPI (blue), MAP2 (green) and α-syn (red) (*a*) or pS129 α-syn (red) (*b, c*). Primary neurons were treated with 100 nM α-syn PFFs at DIV8 for 18 days (*b*) and 22 days (*c*).

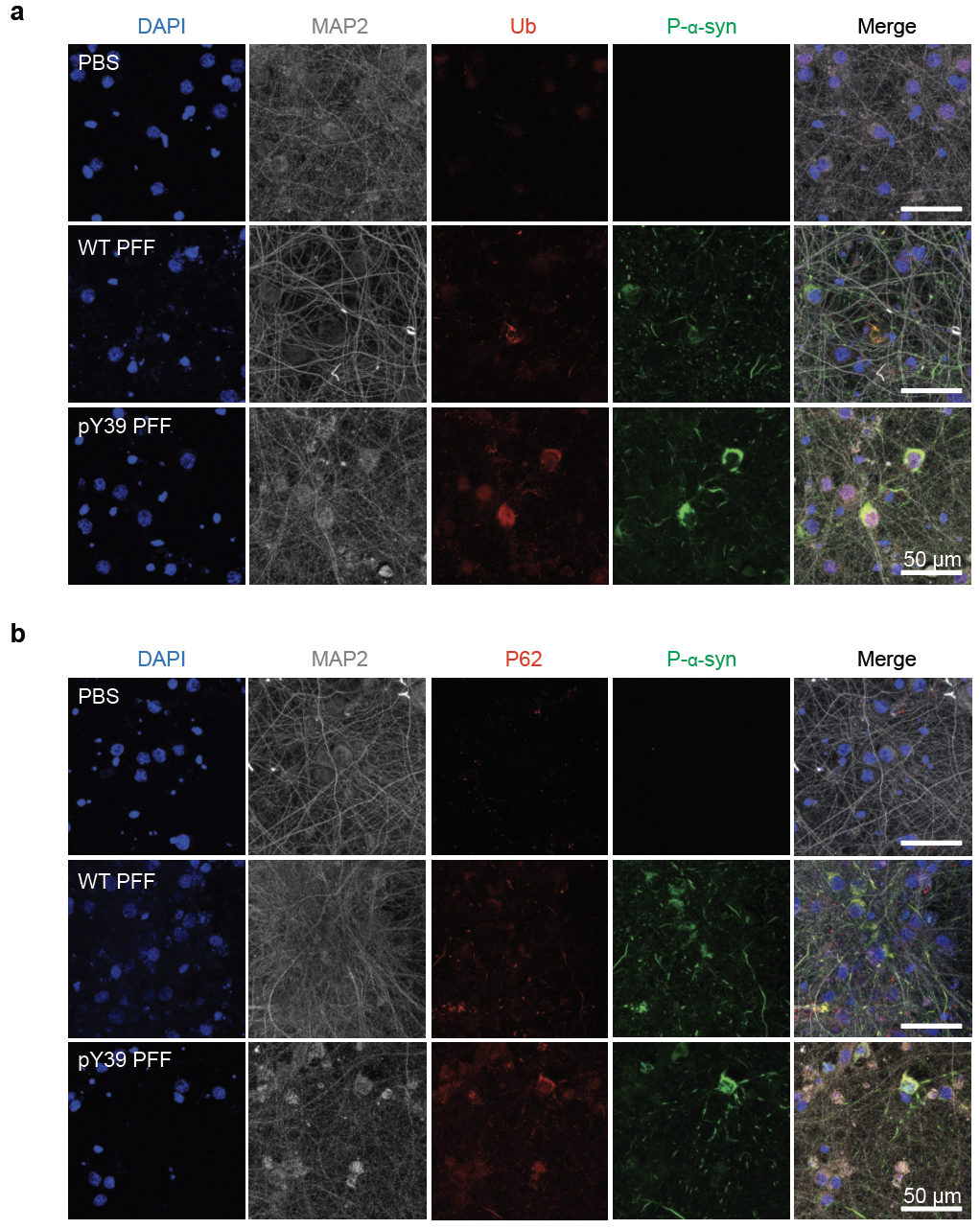

**Figure S6. Co-localization of pY39 PFF-induced pS129 α-syn aggregation with Lewy body markers: ubiquitin (*a*) and P62 (*b*).** Primary neurons were treated with 100 nM α-syn PFFs at DIV8 for 22 days. Confocal microscopic images of fixed neurons stained for DAPI (blue), MAP2 (gray), Ub/P62 (red), and pS129 α-syn (green).

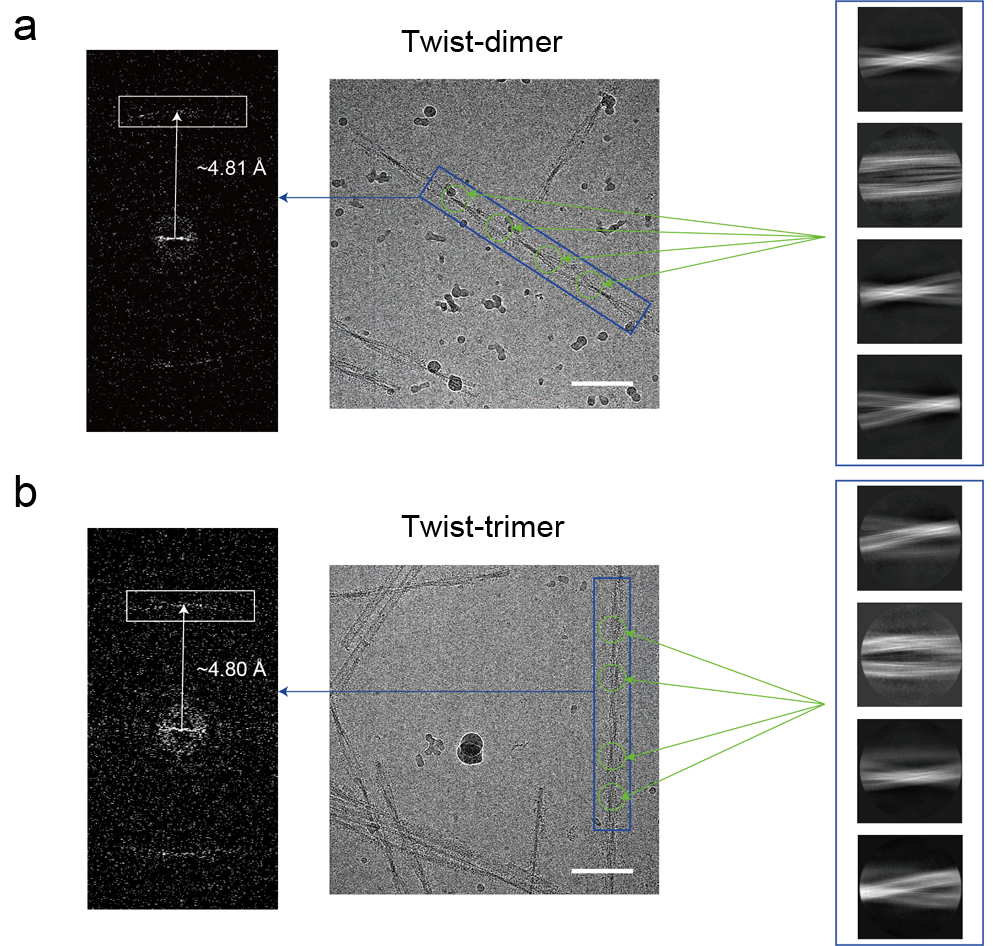

**Figure S7. 2D classification of twist-dimer (*a*) and twist-trimer (*b*).** The power spectra of indicated fibrils are shown on the left. The layer line (pointed by arrow) corresponds to the helical rise. Cryo-EM micrographs are shown in the middle. Scale bar = 100 nm. 2D class averages are shown on the right.

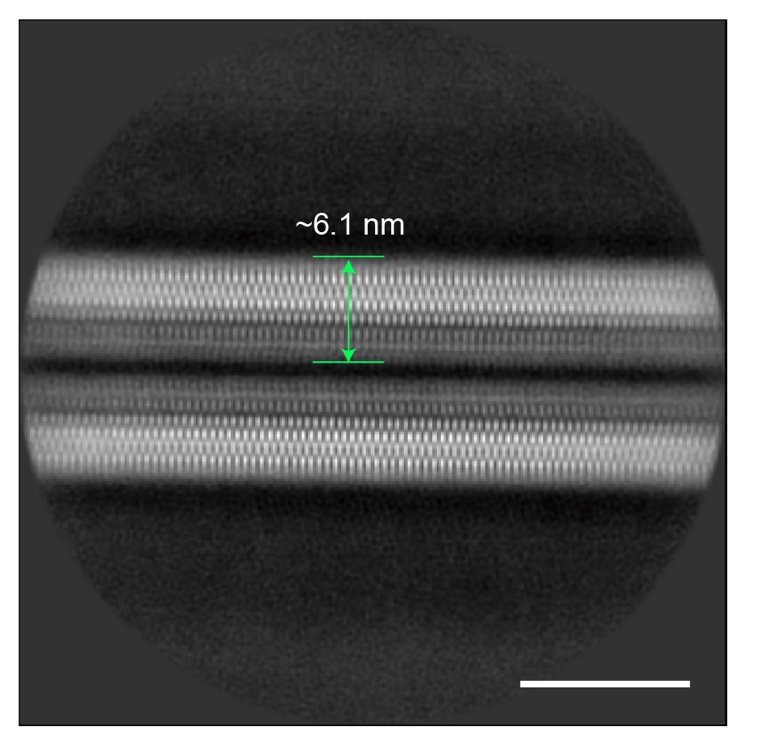

**Figure S8. 2D class averages of pY39 straight fibrils.** Scale bar = 10 nm.

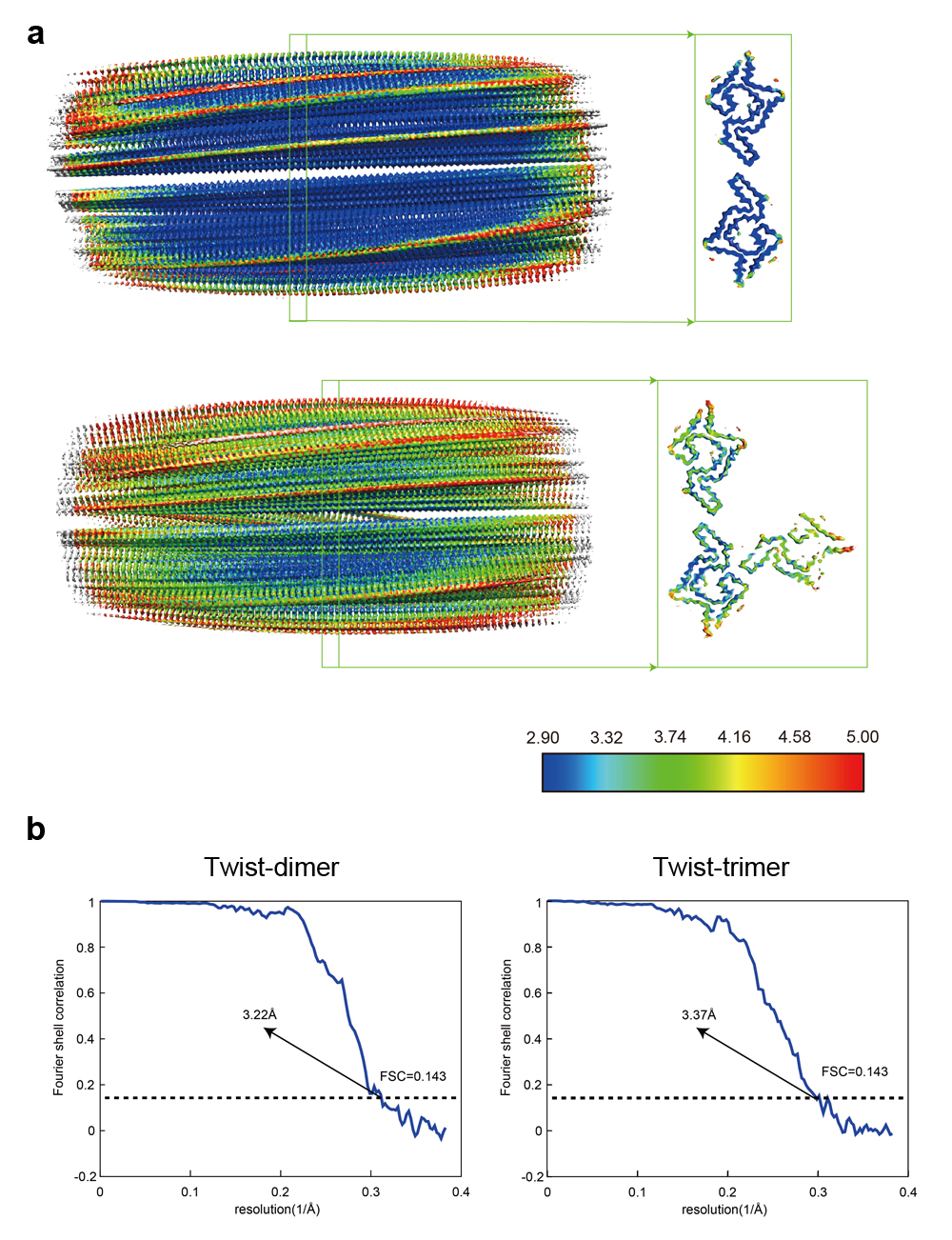

**Figure S9. Resolution estimation of the cryo-EM structures of pY39 α-syn polymorphs.** (*a*) Local resolution estimation. EM reconstruction maps are colored based on the local resolutions of pY39 twist-dimer and twist-trimer fibrils. The color scale indicates the resolution ranging from 2.9 Å to 5.0 Å. (*b*) Gold-standard Fourier shell correlation curve of pY39 twist-dimer and twist-trimer fibrils. The overall resolution of twist-dimer is 3.22 Å and twist -trimer is 3.37 Å.

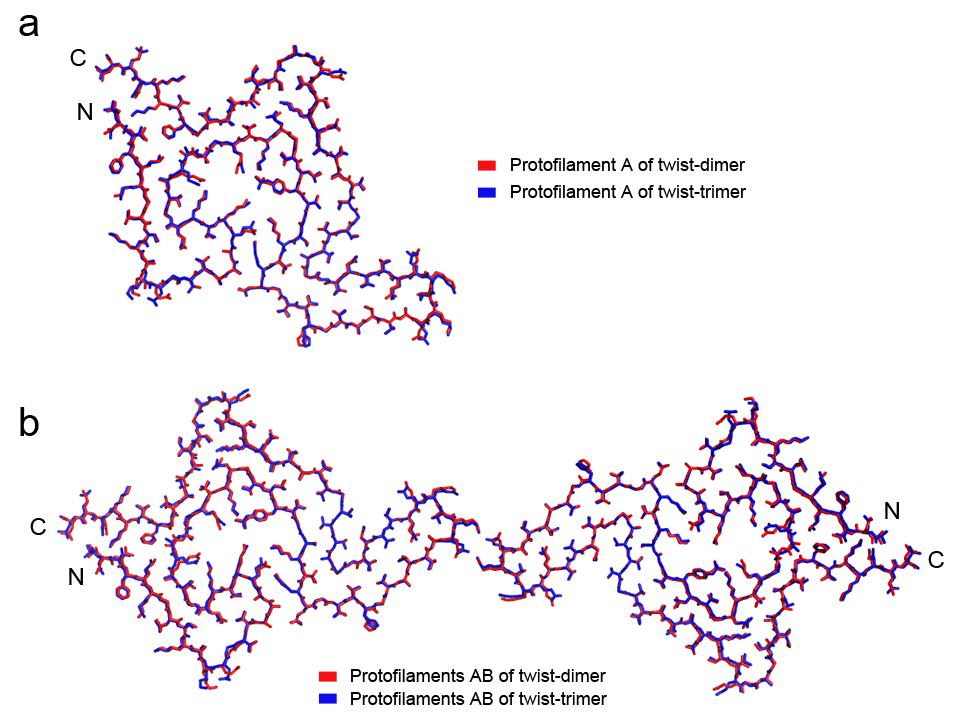

**Figure S10. Structural alignments of pY39 α-syn fibrils.** (*a*) Overlay of α-syn structures in protofilaments A of the twist-dimer (colored in red) and twist-trimer (colored in blue) fibrils. R.m.s.d. of Cα is 0.336 Å. (*b*) Overlay of α-syn dimer structures in protofilaments A and B from the twist-dimer (red) and twist-trimer (blue) fibrils. R.m.s.d. of Cα is 0.406 Å.

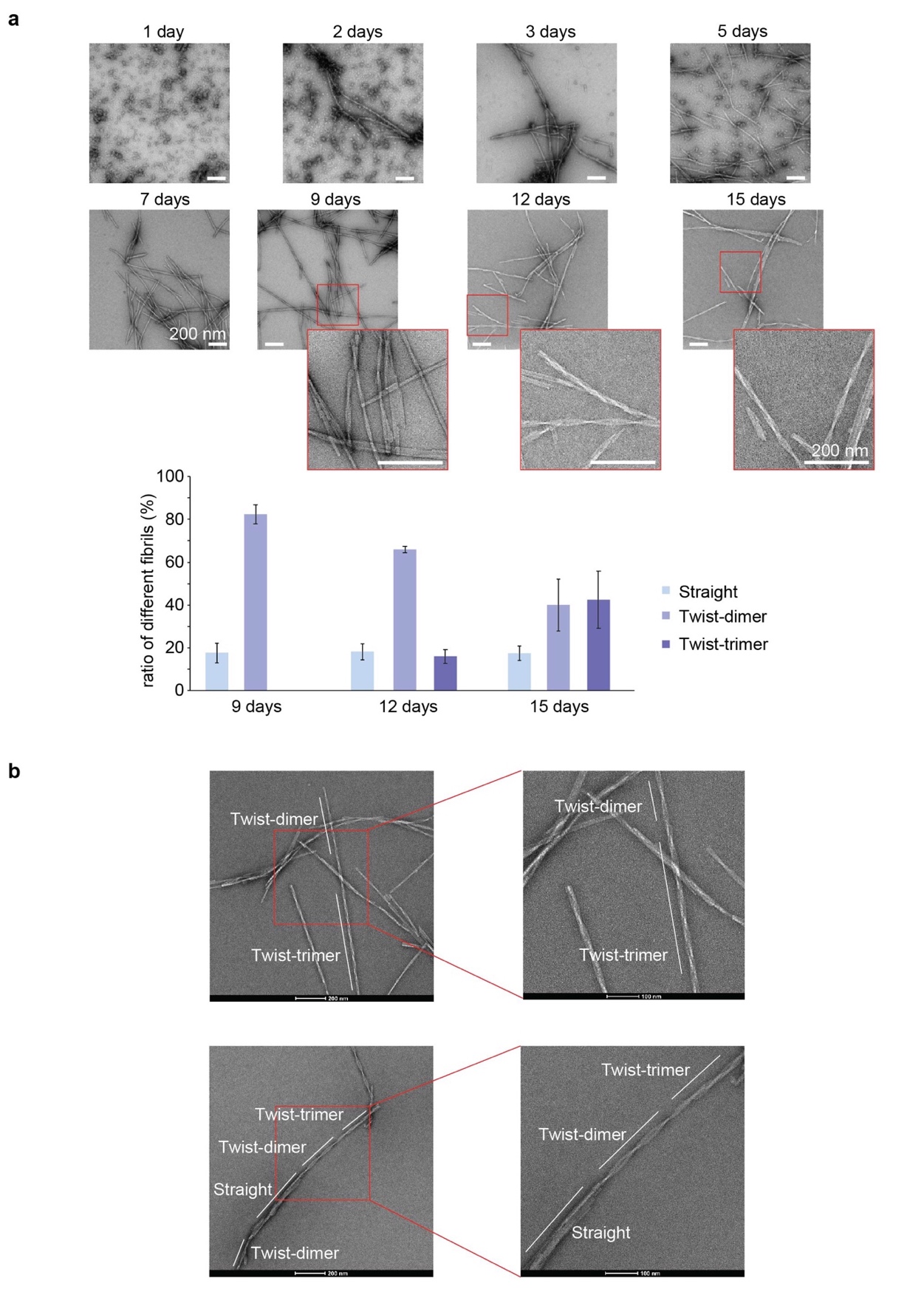

**Figure S11. Successive growth of pY39 α-syn polymorphic fibrils by TEM.** (*a*) Time-course tracking of the fibril formation of pY39 α-syn by TEM. Zoom-in views are shown for the images of 9 days, 12 days and 15 days. Statistical analysis of pY39 polymorphs is shown at the bottom. The percentage of each polymorph is calculated as the fibril length of the polymorph over the total fibril length in the image. Data shown are mean ± s.d., n=3. (*b*) TEM images showing obvious coexistence of different polymorphs in the same fibril.

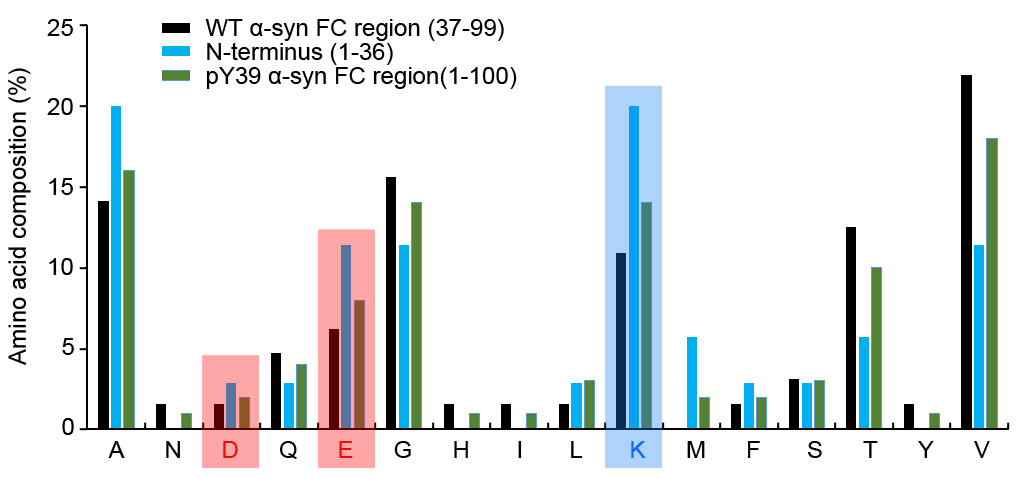

**Figure S12. Amino acid composition of α-syn fragments.** Acidic residues are highlighted in red. Alkaline residues are highlighted in blue. FC: fibril core.

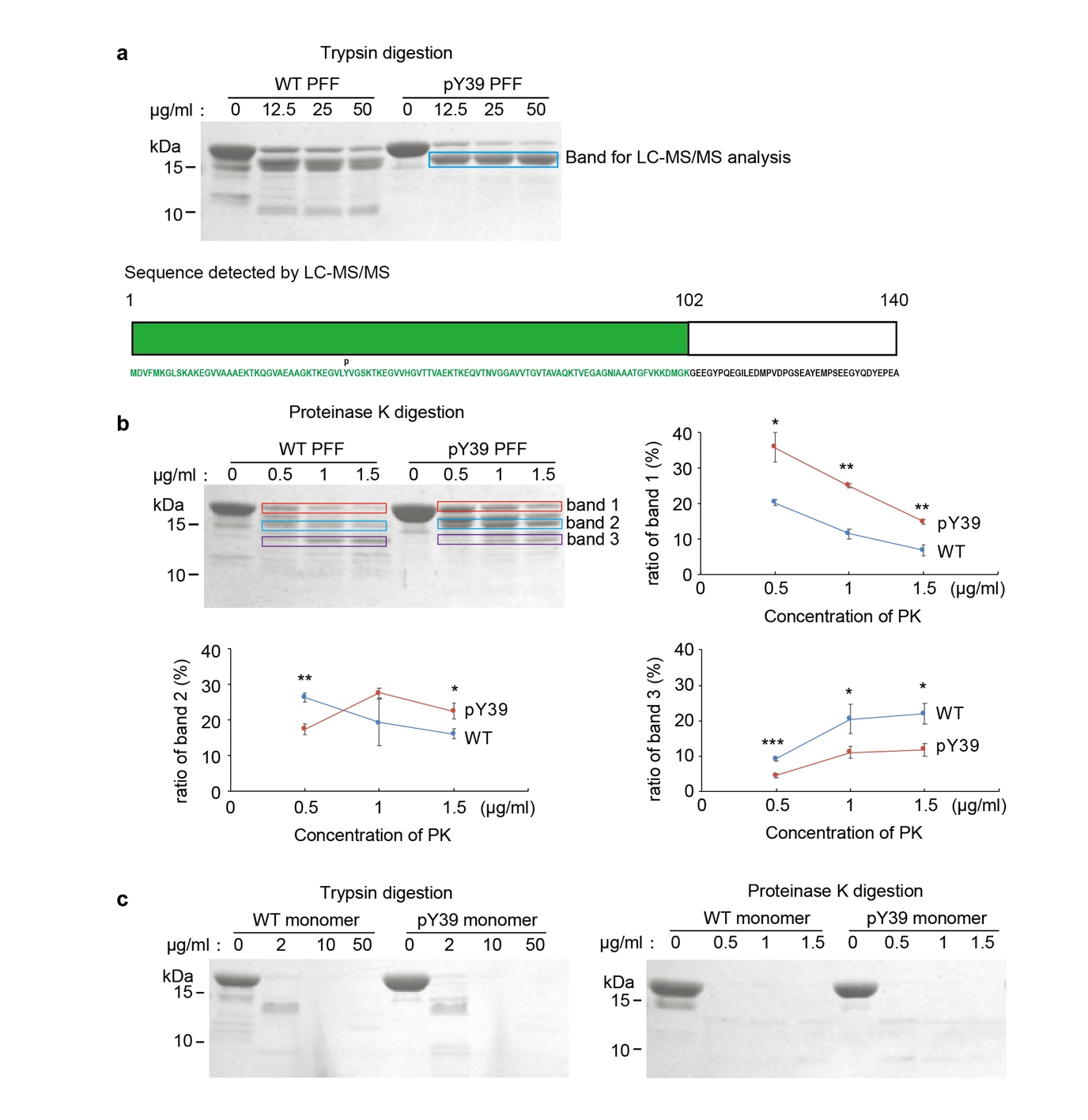

**Figure S13.** **Protease digestion of pY39 and WT α-syn fibrils.**

(*a*) SDS-PAGE of α-syn WT and pY39 PFFs after digestion with trypsin at 37 ºC for 30 min (top). Sequence detected by LC-MS/MS of the indicated band is shown at the bottom. (*b*) SDS-PAGE of α-syn WT and pY39 PFFs after digestion with proteinase K at 37 ºC for 30 min. The intensities of three major bands of the WT and pY39 PFFs after digestion were quantified and normalized to that of the untreated PFFs. Data shown are mean ± s.d., n=3 individual gels. *, *P*<0.05; ** *P*<0.01; *** *P*<0.001. *P* values are based on two-sided Student’s t-test. (*c*) SDS-PAGE of α-syn WT and pY39 monomers.

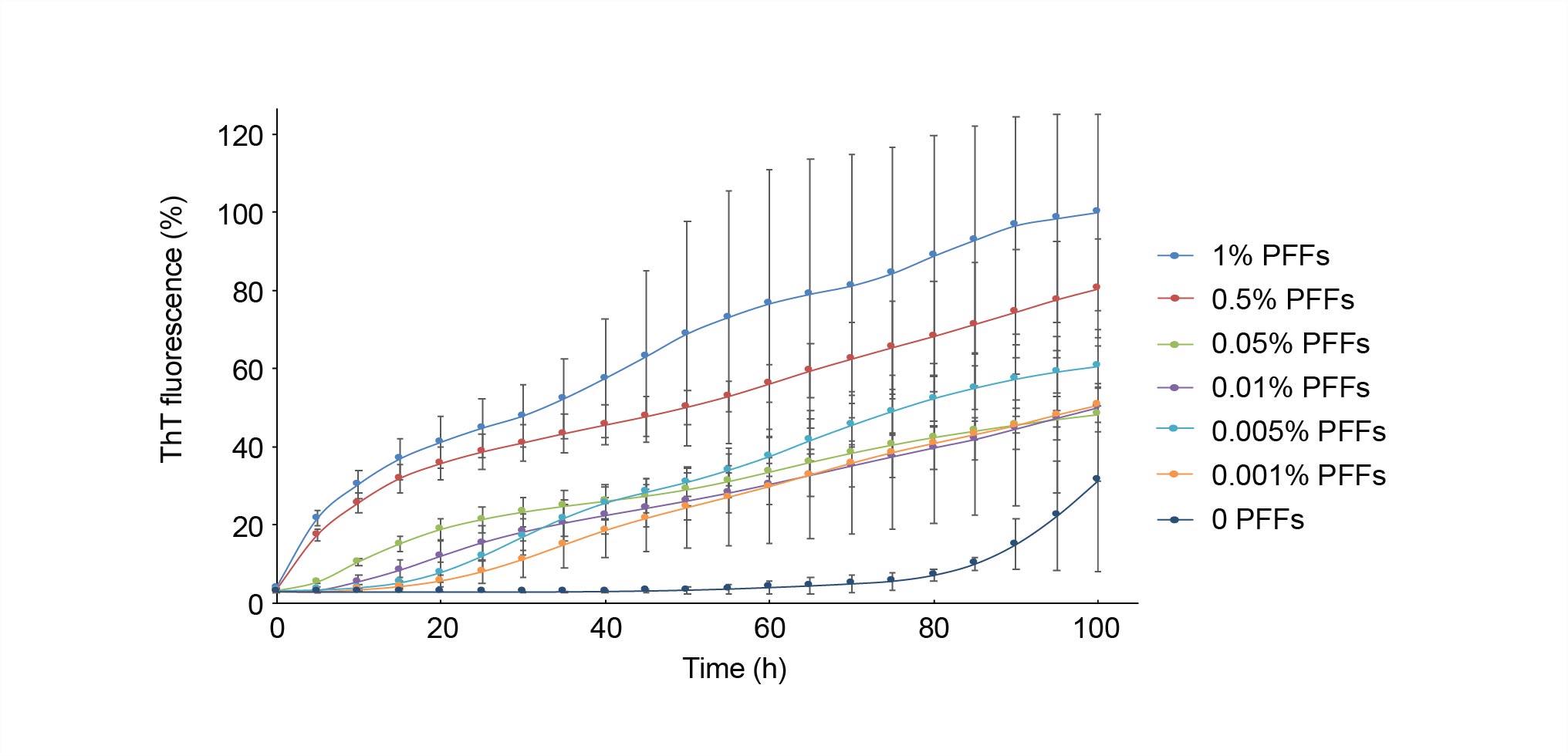

**Figure S14. ThT kinetic assay of WT α-syn aggregation seeded by a gradient percentage (mol%) of WT PFFs.**

Data shown are mean ± s.d., n=3.
